# Accounting for diverse evolutionary forces reveals the mosaic nature of selection on genomic regions associated with human preterm birth

**DOI:** 10.1101/816827

**Authors:** Abigail L. LaBella, Abin Abraham, Yakov Pichkar, Sarah L. Fong, Ge Zhang, Louis J. Muglia, Patrick Abbot, Antonis Rokas, John A. Capra

## Abstract

Human pregnancy requires the coordinated function of multiple tissues in both mother and fetus and has evolved in concert with major human adaptations. As a result, pregnancy-associated phenotypes and related disorders are genetically complex and have likely been sculpted by diverse evolutionary forces. However, there is no framework to comprehensively evaluate how these traits evolved or to explore the relationship of evolutionary signatures on trait-associated genetic variants to molecular function. Here we develop an approach to test for signatures of diverse evolutionary forces, including multiple types of selection, and apply it to genomic regions associated with spontaneous preterm birth (sPTB), a complex disorder of global health concern. We find that sPTB-associated regions harbor diverse evolutionary signatures including evolutionary sequence conservation (consistent with the action of negative selection), excess population differentiation (local adaptation), accelerated evolution (positive selection), and balanced polymorphism (balancing selection). Furthermore, these genomic regions show diverse functional characteristics which enables us to use evolutionary and molecular lines of evidence to develop hypotheses about how these genomic regions contribute to sPTB risk. In summary, we introduce an approach for inferring the spectrum of evolutionary forces acting on genomic regions associated with complex disorders. When applied to sPTB-associated genomic regions, this approach both improves our understanding of the potential roles of these regions in pathology and illuminates the mosaic nature of evolutionary forces acting on genomic regions associated with sPTB.

## INTRODUCTION

Mammalian pregnancy requires the coordination of multiple maternal and fetal tissues^1,2^ and extensive modulation of the maternal immune system so that the genetically distinct fetus is not immunologically rejected^3^. Given this context, pregnancy-related phenotypes and disorders are likely to have experienced diverse selective pressures. This is particularly likely on the human lineage where pregnancy has been shaped by unique human adaptations such as bipedality and enlarged brain size ^4–8^. One major disorder of pregnancy is preterm birth (PTB), a complex multifactorial syndrome^9^ that affects 10% of pregnancies in the United States and more than 15 million pregnancies worldwide each year^10,11^. PTB leads to increased infant mortality rates and significant short- and long-term morbidity^11–14^. Risk for PTB varies substantially with race, environment, comorbidities, and genetic factors^15^. PTB is broadly classified into iatrogenic PTB, when it is associated with medical conditions such as preeclampsia (PE) or intrauterine growth restriction (IUGR), and spontaneous PTB (sPTB), which occurs in the absence of preexisting medical conditions or is initiated by preterm premature rupture of membranes^16–19^. The biological pathways contributing to sPTB remain poorly understood^9^, but diverse lines of evidence suggest that maternal genetic variation is an important contributor^20–24^.

The complexity of human pregnancy and association with unique human adaptations raise the hypothesis that genetic variants associated with birth timing and sPTB have been shaped by diverse evolutionary forces. Consistent with this hypothesis, several immune genes involved in pregnancy have signatures of recent purifying selection^25^ while others have signatures of balancing selection^25–27^. In addition, both birth timing and sPTB risk vary across human populations^28^, which suggests that genetic variants associated with these traits may also exhibit population-specific differences. Variants at the progesterone receptor locus associated with sPTB in the East Asian population show evidence of population-specific differentiation driven by positive and balancing selection^29,30^. Since progesterone has been extensively investigated for sPTB prevention^31^, these evolutionary insights may have important clinical implications. Although these studies have considerably advanced our understanding of how evolutionary forces have sculpted specific genes involved in human birth timing, we lack a comprehensive examination of how diverse evolutionary forces have influenced genomic regions involved in sPTB.

The recent availability of sPTB-associated genomic regions from large genome-wide association studies (GWAS)^32^ coupled with advances in measuring evidence for diverse evolutionary forces—including balancing selection^33^, positive selection^34^, and purifying selection^35^ from human population genomic variation data—present the opportunity to comprehensively survey how evolution has shaped sPTB-associated genomic regions. To achieve this, we developed an approach that identifies evolutionary forces that have acted on genomic regions associated with a complex trait and compares them to appropriately matched control regions. Our approach innovates on current methods by evaluating the impact of multiple different evolutionary forces on trait-associated genomic regions while accounting for genomic architecture-based differences in the expected distribution for each of the evolutionary measures. Application of our approach to 215 sPTB-associated genomic regions showed significant evidence for at least one evolutionary force in 120 regions. Furthermore, we identified functional links to sPTB and other pregnancy phenotypes for representative genomic regions exhibiting evidence for each evolutionary force. These results suggest that a mosaic of evolutionary forces likely influenced human birth timing, and that evolutionary analysis can assist in interpreting the role of specific genomic regions in disease phenotypes.

## RESULTS & DISCUSSION

### Evaluating the significance of evolutionary measures by accounting for genomic architecture

In this study, we compute diverse evolutionary measures on sPTB-associated genomic regions to infer the action of multiple evolutionary forces (Table 1). While various methods to detect signatures of evolutionary forces exist, many of them lack approaches for determining statistically significant observations or rely on the comparison to the distribution of the measure when applied genome-wide^36,37^. Furthermore, population level attributes, such as minor allele frequency (MAF) and linkage disequilibrium (LD), influence the power to detect both evolutionary signatures^38–40^ and GWAS associations^41^. Thus, interpretation and comparison of different evolutionary measures is challenging, especially when the regions under study do not reflect the genome-wide background.

**Table 1:** Evolutionary measures computed on sPTB-associated genomic regions with the corresponding evolutionary signature used to infer the evolutionary force and the associated timescale. GERP: Genomic evolutionary rate profiling. iHS: integrated haplotype score. XP-EHH: cross-population extended haplotype homozygosity (EHH). iES: integrated site-specific EHH. TMRCA: time to most recent common ancestor derived from ARGweaver. Alignment block age was calculated using 100-way multiple sequence alignments to determine the oldest most recent common ancestor for each alignment block.

| Measures | Evolutionary signature | Evolutionary force | Time scale |
| --- | --- | --- | --- |
| PhyloP | Substitution rate | Positive/negative selection | Across species |
| PhastCons | Sequence conservation | Negative selection | Across species |
| GERP |  |  |  |
| LINSIGHT |  |  | Across species and human populations |
| F<sub>ST</sub> | Population differentiation | Local adaptation | Human populations |
| iHS | Haplotype homozygosity | Positive selection | Human populations |
| XP-EHH |  |  |  |
| iES |  |  |  |
| Beta Score | Balanced polymorphisms | Balancing selection | Human populations |
| Allele Age (TMRCA) | Ancestral recombination graphs/Alignments | Evolutionary origin / Negative selection | Human populations |
| Alignment block age | Sequence conservation | Evolutionary origin / Negative selection | Across species |

Here we develop an approach that derives a matched null distribution accounting for MAF and LD for each evolutionary measure and set of regions. We generate 5,000 control region sets that each match the trait-associated regions on these attributes (Methods). Then, to calculate an empirical p-value and z-score for each evolutionary measure and region of interest, we compare the median values of the evolutionary measure for variants in the sPTB-associated genomic region to the same number of variants in the corresponding matched control regions. This enables comparison across evolutionary measures and genomic regions (Figure 1A, Methods).

**Figure 1:**
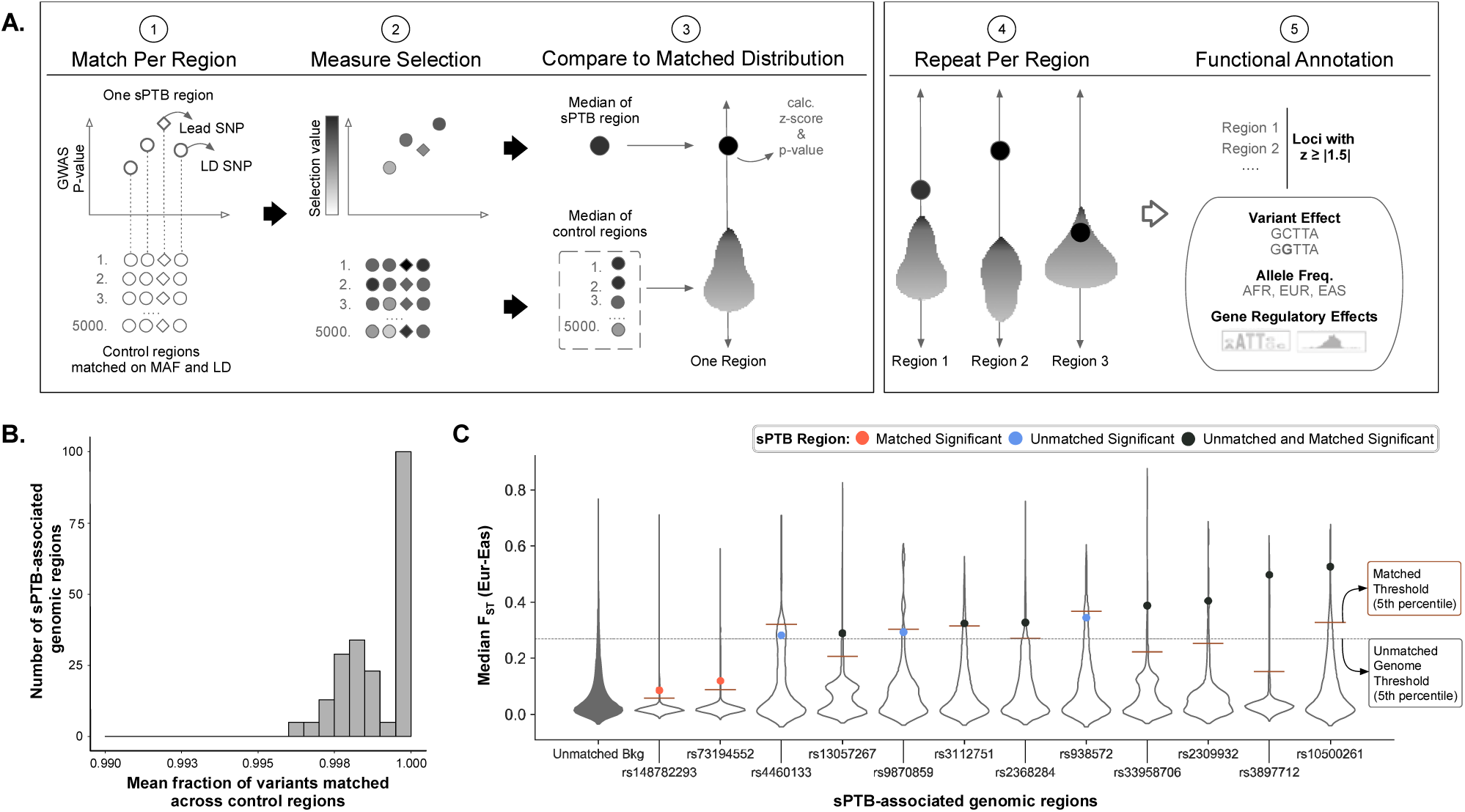
Accounting for minor allele frequency and linkage disequilibrium of sPTB-associated genomic regions identifies loci that have experienced diverse evolutionary forces. **A.** We compared evolutionary measures for each sPTB-associated genomic region (n=215) to MAF and LD matched control regions (∼5000). The sPTB-associated genomic regions each consisted of a lead variant (p<10E-4 association with sPTB) and variants in high LD (r^2^>0.9) with the lead variant. Each control region has an equal number of variants as the corresponding sPTB-associated genomic region and is matched for MAF and LD (‘Match Per Region’). We next obtained the values of an evolutionary measure for the variants included in the sPTB-associated regions and all control regions (‘Measure Selection’). The median value of the evolutionary measure across variants in the sPTB-associated region and all control regions was used to derive a z-score (‘Compare to Null Distributions’). We repeated these steps for each sPTB-associated region (‘Repeat Per Region’) and then functionally annotated sPTB-associated regions with absolute z-scores ≥ 1.5 (‘Functional Annotation’). **B.** Across all sPTB-associated genomic regions, the mean fraction of variants matched across all control regions was ≥ 0.99. **C.** A representative example for pairwise F_ST_ between East Asians (EAS) and Europeans (EUR). Violin plots display the unmatched genome background (‘unmatched Bkg’, dark fill) or the matched background (no fill) with the sPTB-region labeled by the lead variant (rsID), i.e. the variant with the lowest sPTB-association p-value. Each point represents the median value of F_ST_ for the sPTB-associated regions labeled by the lead SNP (rsID, x-axis). The dotted line represents the threshold for the top 5^th^ percentile when randomly sampled (n=5,000) from the ‘unmatched Bkg.’ The solid horizontal line for each unfilled violin plot represents the 5^th^ percentile for matched background distribution. Each dot’s color represents whether the region is significant by genome background (blue), matched background (red), or both (black). Note that significance is influenced by choice of background and that our approach identifies sPTB-associated regions that have experienced diverse evolutionary forces while controlling for important factors, such as MAF and LD.

We used this approach to evaluate the evolutionary forces acting on genomic regions associated with sPTB. First, we identified all variants nominally associated with sPTB (p<10E-4) in the largest available GWAS^32^ and grouped variants into regions based on high LD (r^2^>0.9). It is likely that many of these nominally associated variants affect sPTB risk, but did not reach genome-wide significance due to factors limiting the statistical power of the GWAS^32^. Therefore, we assume that many of the variants with sPTB-associations below this nominal threshold contribute to the genetic basis of this trait. This identified 215 independent sPTB-associated genomic regions, which we refer to by the lead variant (SNP or indel with the lowest p-value in that region).

For each of the 215 sPTB-associated genomic regions, we generated control regions as described above. The match quality per genomic region, defined as the fraction of sPTB variants with a matched variant averaged across all control regions, is ≥ 99.6% for all sPTB-associated genomic regions (Figure 1B). The matched null distribution aggregated from the control regions varied substantially between sPTB-associated genomic regions for each evolutionary measure and compared to the unmatched genome-wide background distribution (Supplemental Figure 1). The F_ST_ measure between East Asians and Europeans (F_ST-EurEas_) illustrates this variation. Different sets of sPTB-associated genomic regions had statistically significant (p<0.05) median F_ST-EurEas_ based on comparison to the unmatched genome-wide distribution versus comparison to the matched null distribution (Figure 1C). Two regions (Figure 1C: red dots) were statistically significant (p<0.05) only when using our method due to the narrow shape of the matched null distribution. In contrast, three other regions (Fig 1C: blue dots) were significant based on the genome-wide distribution, but not using our method likely due to the genetic architecture of this region. Seven regions (Fig 1C: black dots) were statistically significant using both methods. Similar results were obtained across the other evolutionary measures (See Supplemental Figure 1 for break down by evolutionary measure).

Our approach to test for signatures of different evolutionary forces has many advantages. Comparing evolutionary measures against a null distribution that accounts for MAF and LD enables us to increase the sensitivity with which we can infer the action of evolutionary forces on sets of genomic regions that differ in their genome architectures. In addition, the lead SNPs assayed in a GWAS are often not the causal variant, so by testing both the lead SNPs and those in LD when evaluating a genomic region for evolutionary signatures, we are able to better represent the trait-associated evolutionary signatures compared to other methods that evaluate only the lead variant^42^ or all variants, including those not associated with the trait, in a genomic window^43^ (Supplementary Table 1). Finally, our approach uses an empirical framework that leverages the strengths of diverse existing evolutionary measures.

### Genomic regions associated with sPTB exhibit signatures of diverse modes of selection

To gain insight into the modes of selection that have acted on sPTB-associated genomic regions, we focused on genomic regions with extreme evolutionary signatures by selecting the 120 sPTB-associated regions with at least one extreme z-score (z ≥ +/- 1.5) (Figure 2; Supplementary Tables 2 and 3) for further analysis. Notably, each evolutionary measure had at least one genomic region with an extreme observation (p<0.05). Hierarchical clustering of the 120 regions revealed 12 clusters of regions with similar evolutionary patterns. We manually combined the 12 clusters based on their dominant evolutionary signatures into five major groups with the following general evolutionary patterns (Figure 2): conservation/negative selection (group A: clusters A1-4), excess population differentiation/local adaptation (group B: clusters B1-2), positive selection (group C: cluster C1), long-term balanced polymorphism/balancing selection (group D: clusters D1-2), and other diverse evolutionary signatures (group E: clusters E1-4).

**Figure 2:**
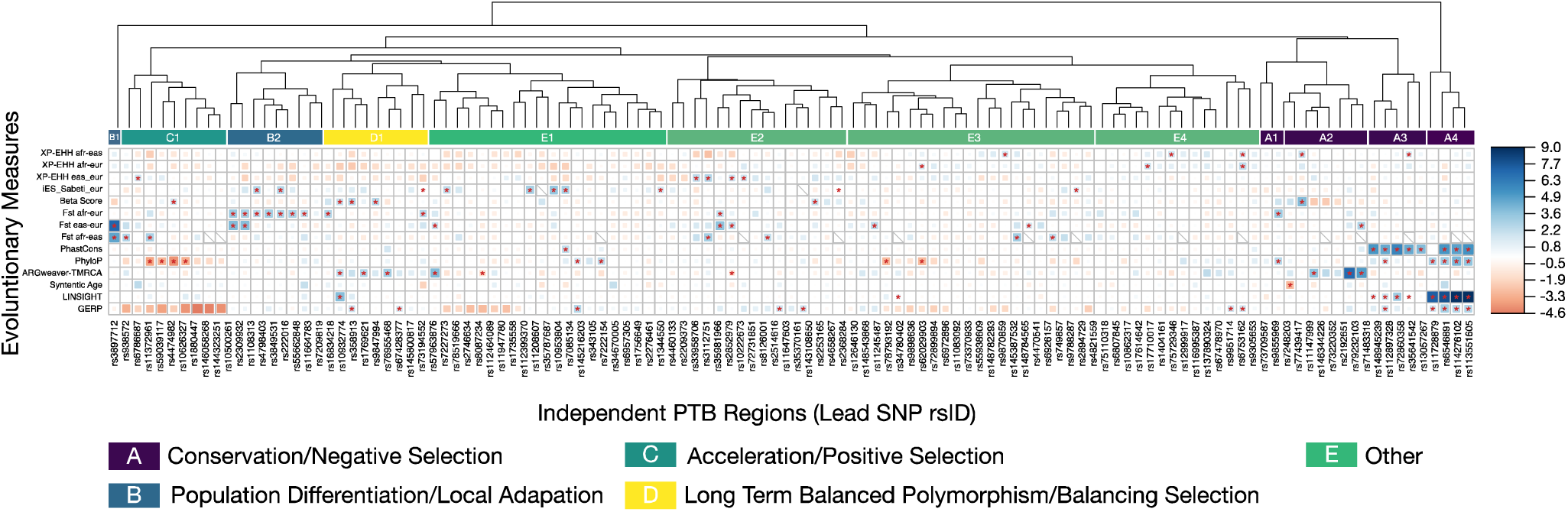
Clusters of sPTB-associated genomic regions have experienced diverse evolutionary forces. We tested sPTB-associated genomic regions (x-axis) for diverse types of selection (y-axis), including F_ST_ (population differentiation), XP-EHH (positive selection), Beta Score (balancing selection), allele age (time to most recent common ancestor, TMRCA, from ARGweaver), alignment block age, phyloP (positive/negative selection), GERP, LINSIGHT, and PhastCons (negative selection) (Table 1). The relative strength (size of square) and direction (color) of each evolutionary measure for each sPTB-associated region is presented as a z-score calculated from that region’s matched background distribution. Only regions with |z| ≥ 1.5 for at least one evolutionary measure before clustering are shown. Statistical significance was assessed by comparing the median value of the evolutionary measure to the matched background distribution to derive an empirical p-value (*p>0.05). Hierarchical clustering of sPTB-associated genomic regions on their z-scores identifies distinct groups or clusters that appear to be driven by different types of evolutionary forces. Specifically, we interpret regions that exhibit higher than expected values for PhastCons, PhyloP, LINSIGHT, and GERP to have experienced conservation and negative selection (Group A); regions that exhibit higher than expected pairwise F_ST_ values to have experienced population differentiation/local adaptation (Group B); regions that exhibit lower than expected values for PhyloP to have experienced acceleration/positive selection (Group C); and regions that exhibit higher than expected Beta Score and older allele ages (TMRCA) to have experienced balancing selection (Group D). The remaining regions exhibit a variety of signatures that are not consistent with a single evolutionary mode (Group E).

Previous literature on complex genetic traits^44–46^ and pregnancy disorders^25,29,30,47,48^ supports the finding that multiple modes of selection have acted on sPTB-associated genomic regions. Unlike many of these previous studies that tested only a single mode of selection, our approach tested multiple modes of selection. Of the 215 genomic regions we tested, 9% had evidence of conservation, 5% had evidence of excess population differentiation, 4% had evidence of accelerated evolution, 4% had evidence of long-term balanced polymorphisms, and 34% had evidence of other combinations. From these data we infer that negative selection, local adaptation, positive selection, and balancing selection have all acted on genomic regions associated with sPTB, highlighting the mosaic nature of the evolutionary forces that have shaped this trait. In addition to differences in evolutionary measures, variants in these groups also exhibited differences in their functional effects, likelihood of influencing transcriptional regulation, frequency distribution between populations, and effects on tissue-specific gene expression (Figure 3; Supplementary Tables 4 and 5). We now describe each group and give examples of their members and their potential connection to PTB and pregnancy.

**Figure 3.**
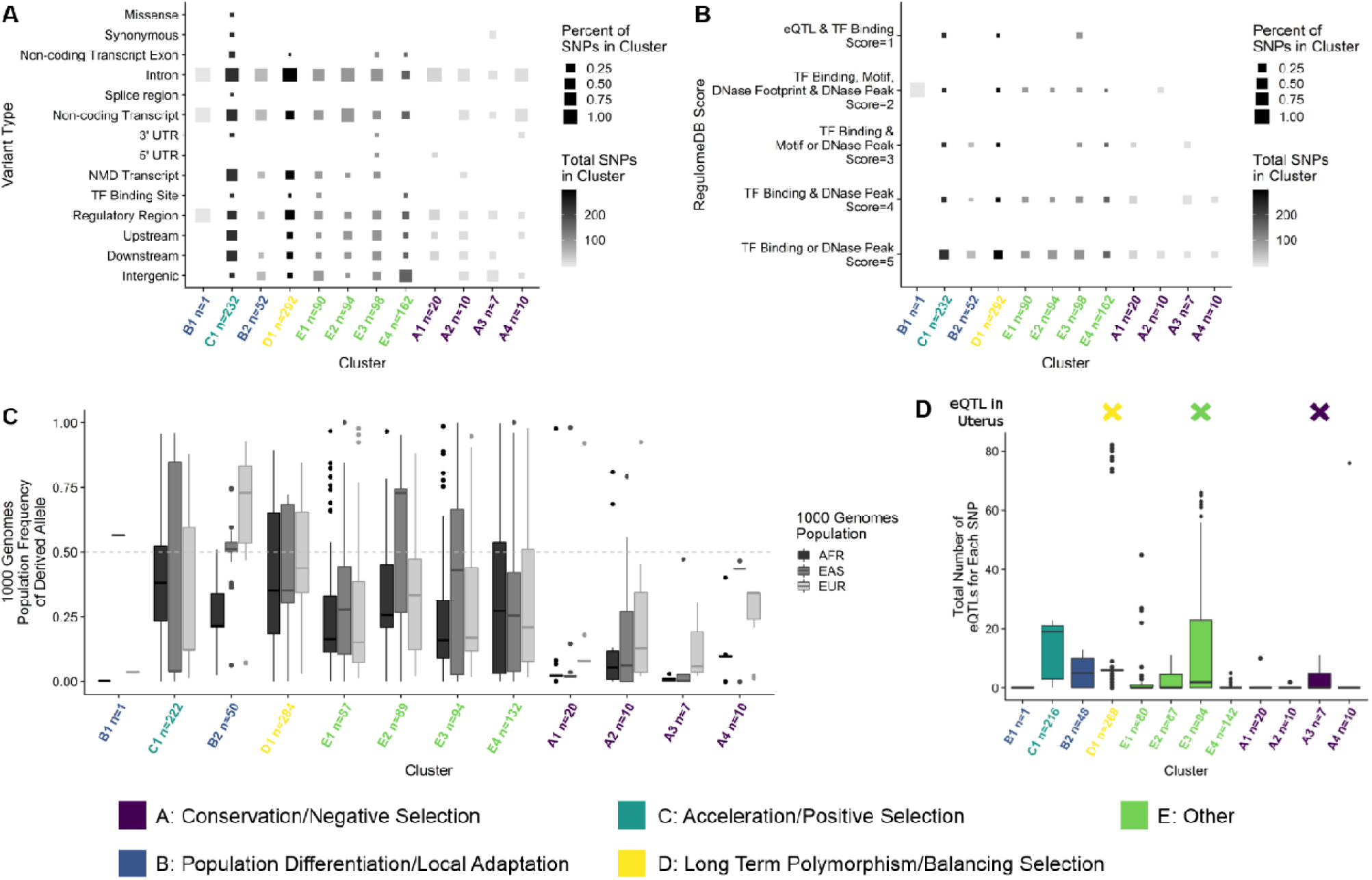
Clusters of PTB regions that have experienced different types of selection vary widely in their molecular characteristics and functions. Clusters are ordered as they appear in the z-score heatmap (Fig. 2) and colored by their major type of selection: Group A: Conservation and negative selection (Purple), Group B: Population differentiation/local adaptation (Blue), Group C: Acceleration and positive selection (Teal), Group D: Long term polymorphism/balancing selection (Teal), and Other (Green). **A.** The proportions of different types of variants (e.g., intronic, intergenic, etc.) within each cluster (x-axis) based on the Variant Effect Predictor (VEP) analysis. Furthermore, cluster C1 exhibits the widest variety of variant types and is the only cluster that contains missense variants. Most variants across most clusters are located in introns. **B.** The proportion of each RegulomeDB score (y-axis) within each cluster (x-axis). Most notably, PTB regions in three clusters (B1, A5, and D4) have variants that are likely to affect transcription factor binding and linked to expression of a gene target (Score=1). Almost all clusters contain some variants that are likely to affect transcription factor binding (Score=2). **C.** The derived allele frequency (y-axis) for all variants in each cluster (x-axis) for the African (AFR), East Asian (EAS), and European (EUR) populations. Population frequency of the derived allele varies within populations from 0 to fixation. **D.** The total number of eQTLs (y-axis) obtained from GTEx for all variants within each cluster (x-axis) All clusters but one (C2 with only one variant) have at least one variant that is associated with the expression of one or more genes in one or more tissues. Clusters A1, A5, and D4 also have one or more variants associated with expression in the uterus.

### Group A: Sequence Conservation/Negative selection

Group A contains 19 genomic regions and 47 total variants. Variants in this group had higher than expected values for evolutionary measures of sequence conservation and alignment block age: PhastCons100, PhyloP, allele age in TMRCA derived from ARGweaver, LINSIGHT and/or GERP (Figure 2). The strong sequence conservation suggests that these genomic regions evolved under negative selection. The average derived allele frequency of group A variants across populations is 0.15 (Figure 3C). The majority of variants are intronic (37/47: 79%) but a considerable fraction is intergenic (8/47: 17%; Figure 3B). One coding variant (rs17436878) results in a synonymous change in the gene *RGL1* and another variant (rs6546891) is located in the 3’ UTR of the Ten-Eleven Translocation Methylcytosine Dioxygenase 3 (*TET3*) gene. Only one variant is predicted to affect regulatory binding (rs71483318, RegulomeDB score=2), a finding consistent with the observation that most (41/47) variants in this group are not known to be associated with expression changes (Figure 4D).

**Figure 4:**
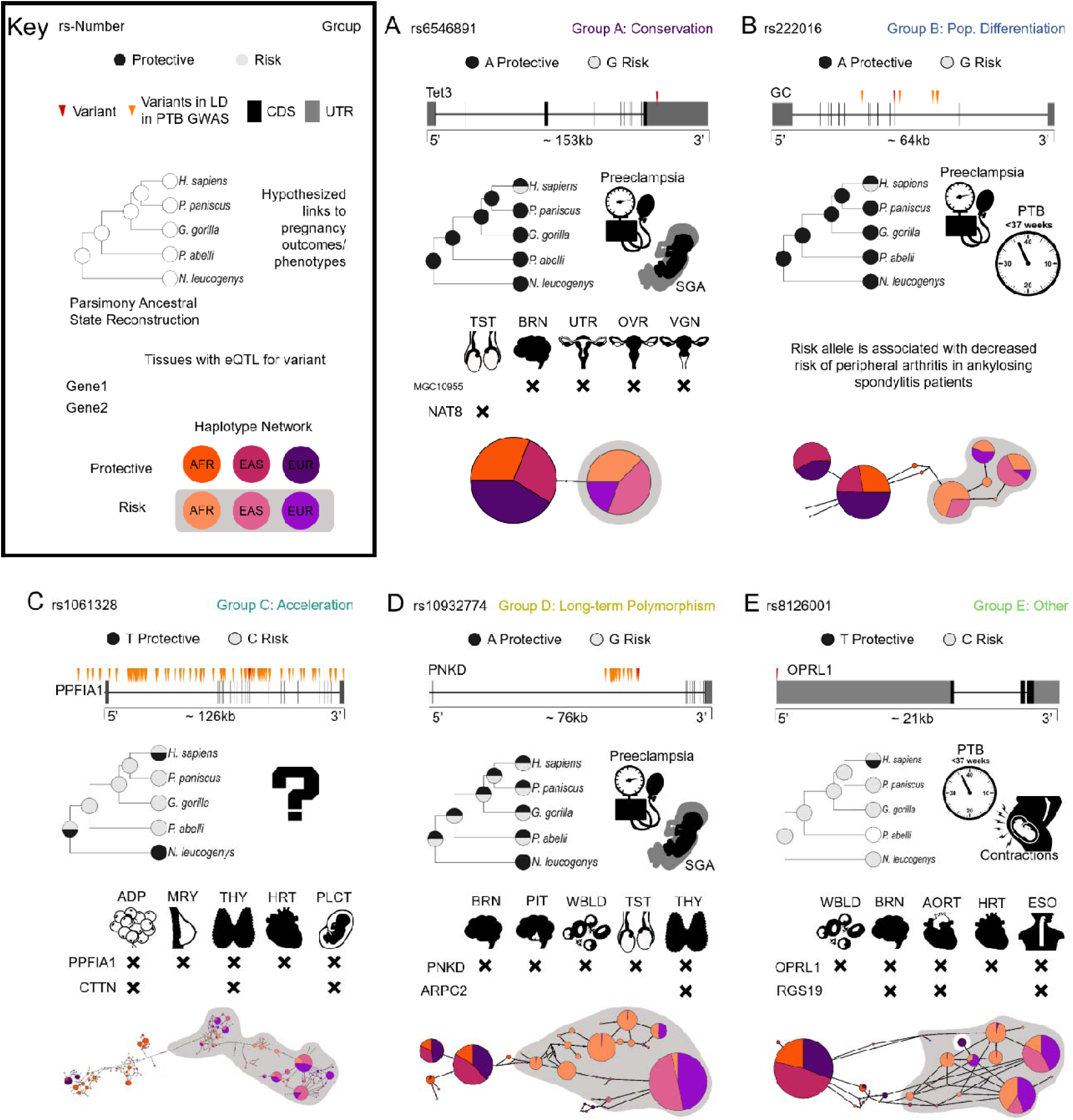
Functional and evolutionary characterizations of a representative variant from each group illustrate their diverse histories and roles in risk of preterm birth. For each variant of interest, we report the following information listed from top to bottom: the protective and risk alleles as predicted by the sPTB GWAS^136^; the location relative to the nearest gene of the variant and linked variants; the allelic data at this location across the great apes and the parsimony reconstruction of the ancestral allele(s) at this site; hypothesized links to pregnancy outcomes or phenotypes from previous studies; selected significant GTEx hits; and finally, the human haplotype(s) containing each variant in a haplotype map labeled by 1KG population. **A.** Representative variant from group A (conservation): rs6546894 contains a human-specific risk allele and is located in the 3’ UTR of the gene *TET3*. The site has strong evidence of long-term evolutionary conservation. The gene *TET3* has been shown to be elevated in preeclamptic and small for gestational age (SGA) placentas^53^. The rs6546894 variant is also associated with changes in expression of genes *MGC10955* and *NAT8* in the testis (TST), brain (BRN), uterus (UTR), ovaries (OVR), and vagina (VGN) tissues. The variant does not have high LD (r^2^ > 0.9) with any other 1KG variants so there are only two haplotypes/alleles. **B.** Representative variant from group B (population differentiation): rs222016 is located in the intron of the gene *GC* and has a human-specific protective allele. The gene *GC* is linked the occurrence of sPTB^65^. This variant is not associated with any known eQTLs, but the risk allele has been associated with a protective effect on arthritis^59^. Haplotypes containing the protective allele are rare in the African population. **C.** Representative variant from group C (acceleration): rs1061328 is located in an intron of the gene *PPFIA1* and is in high LD with 156 other variants spanning the gene’s length. It has a protective allele that is human-specific. This variant is associated with changes in expression of *PPFIA1* and *CTTN* genes in adipose cells (ADP), mammary tissue (MRY), the thyroid (THY), and heart (HRT). The gene *CTTN* has been shown to be expressed in placental cells^85,86^. There is a total of 102 haplotypes associated with this variant. **D.** Representative variant from group D (long-term polymorphism): rs10932774 is located in the intronic region of the gene *PNKD* and is in high LD with 27 additional SNPs in an 8.8 kb region. Consistent with the action of balancing selection, both alleles of the variant are found throughout the great apes. The gene *PNKD* is upregulated in severely preeclamptic placentas^99^ and *ARPC2* has been associated with SGA^137^. Expression changes associated with this variant include *PNKD* and *ARPC2* in the brain, pituitary gland (PIT), whole blood (WBLD), testis, and thyroid (THY). Haplotypes containing the risk allele are more prevalent across populations and also display greater haplotype diversity. **E.** Representative variant from group E (other): rs8126001 is located in the 5’ UTR of the gene *OPRL1* and has a human-specific protective allele. The protein product of the *ORPL1* gene is the nociceptin receptor, which has been linked to contractions and the presence of nociception in preterm uterus samples^115,116^. This variant is associated with expression of the genes *OPRL1* and *RGS19* in whole blood, the brain, aorta (AORT), heart, and esophagus (ESO: eQTL data is from GTEx v6). There is more haplotype diversity in the risk allele and these haplotypes are more prevalent in the European and African populations.

The sPTB-associated genomic region in the 3’ UTR of the gene *TET3* has significant values for LINSIGHT, GERP, and PhastCons100 (Figure 2). The G allele of rs6546891 is associated with a nominally increased risk of sPTB in European ancestry individuals (OR: 1.13; adjusted p-value: 5.4×10^−5^)^32^. This risk allele (G) arose on the human lineage while the protective allele (A) is ancestral and is present across the great apes (Figure 4A). The risk allele is the minor allele in all three populations examined and has the lowest frequency in Europeans. Functionally, this variant is an eQTL for 76 gene/tissue pairs: most notably this variant is associated with the expression of the hypothetical protein LOC730240 in the uterus, ovary, vagina, and brain as well as the expression of N-acetyltransferase 8 (*NAT8*) in the testis. The conservation detected at this locus suggests that disruptions in *NAT8* or *TET3* are likely to be deleterious.

The genes *TET3* and *NAT8* are both linked to gestation and pregnancy outcomes. In mice, *TET3* affects epigenetic reprogramming in oocytes and zygotes, is required for neonatal growth, and depletion of *TET3* in female mice results in reduced fecundity^49,50^. In humans, *TET3* expression was detected in the villus cytotrophoblast cells in the first trimester as well as in maternal decidua of placentas^51^. Expression profiling showed elevated *TET3* transcripts in preeclamptic placentas and in pregnancies ending with the birth of a newborn that is small for gestational age (SGA)^52^. *TET3* is also hypothesized to play a role in the link between preterm birth and the risk of neurodevelopmental disorders due to the gene’s role in epigenetic regulation^53^. *NAT8*, which is involved in acetylation of histones, may also play a role in epigenetic changes during pregnancy^54^. The strong sequence conservation of the region containing variant rs6546891 is consistent with previous findings that negative selection is the dominant mode of selection for *TET3*^55^. More broadly, these findings suggest that several sPTB-associated genomic regions have experienced negative selection, consistent with previous studies^42,56^.

### Group B: Population Differentiation/Population-specific Adaptation

Group B (clusters B1 and B2) contained variants with a higher than expected differentiation (F_ST_) between pairs of human populations (Figure 2). There were 10 sPTB-associated genomic regions in this group, which contain 53 variants. Most variants in this group are intronic (38/53) while the rest are intergenic (14/53) or located within 5000bp upstream of a gene (1/53) (Figure 3A). One variant (rs3897712) may be involved in regulating transcription factor binding (Figure 3B), but is not a known eQTL. For the remaining variants, the majority are an eQTL in at least one tissue (29/52; Figure 3D). The derived allele frequency in cluster B1 is high in East Asian populations and very low in African and European populations (Figure 3C). We found that 3 of the 10 lead variants have higher risk allele frequencies in African compared to European or East Asian populations. This is noteworthy because the rate of PTB is twice as high among black women compared to white women in the United States^57,58^. These three variants are associated with expression levels of the genes *SLC33A1, LOC645355*, and *GC*, respectively.

The six variants within the sPTB-associated region near *GC*, Vitamin D Binding Protein, are of particular interest. The lead variant is rs222016. The G allele of this lead variant (rs222016) has a higher frequency in African populations, is the ancestral allele, and is associated with increased risk of sPTB (European cohort, OR: 1.15; adjusted p-value 3.58×10-5; Figure 4B)^32^. This variant has also been associated with vitamin D levels and several other disorders; for example, in individuals with ankylosing spondylitis the G allele (risk for sPTB) is associated with increased risk of developing peripheral arthritis^59^. This variant is also associated with high baseline D3 levels in serum but is not associated with reduced risk of D3 insufficiency^60^. There is also evidence that vitamin D levels prior to delivery are associated with sPTB,^61–64^ that levels of GC in cervico-vaginal fluid may help predict sPTB^65,66^, and that vitamin D deficiency may contribute to racial disparities in birth outcomes^67,68^. Specifically, vitamin D deficiency may contribute to potential risk for preeclampsia among Hispanic and African American women^69^. The population-specific differentiation associated with the variant rs222016 is consistent with the differential evolution of the vitamin D system between populations—likely in response to different environments and associated changes in skin pigmentation^70,71^. Our results provide evolutionary context for the link between vitamin D and pregnancy outcomes^72^ and suggest a role for variation in the gene *GC* in the ethnic disparities in pregnancy outcomes.

### Group C: Accelerated substitution rates/Positive selection

Variants in cluster C1 (group C) had lower than expected values of PhyloP. This group contains nine sPTB-associated genomic regions and 232 total variants. The large number of linked variants is consistent with the accumulation of polymorphisms in regions undergoing positive selection. The derived alleles in this group show no obvious pattern in allele frequency between populations (Figure 3C). While most variants in this group are intronic (218/232), there are missense variants in the genes Protein Tyrosine Phosphatase Receptor Type F Polypeptide Interacting Protein Alpha 1 (*PPFIA1)* and Plakophilin 1 (*PKP1;* Figure 3A). Additionally, 16 variants in this group are likely to affect transcription factor binding (regulomeDB score of 1 or 2; Figure 3B). Consistent with this finding, of the 216 variants tested in GTEx, 167 are associated with expression of at least one gene in one tissue (Figure 3C).

The lead variant associated with *PPFIA1* (rs1061328) is linked to an additional 156 variants, which are associated with the expression of a total of 2,844 tissue/gene combinations. This variant (rs1061328) has signatures of positive selection and is associated with the expression of genes involved in cell adhesion and migration—critical processes in the development of the placenta. There are three alleles at this locus—two (C and T) were examined in the sPTB GWAS, while the third (G allele) is rare. The risk allele, C, is ancestral and is the major allele in the European and East Asian populations. The derived protective allele (effect: 0.868, adjusted p-value 5.05×10^−4^)^32^ is the major allele in the African population. The C→T polymorphism is synonymous (GAC→CAT) at an asparagine residue in the 14th of 28 exons in the *PPFIA1* gene. A third derived allele (G) is present at very low frequency (<0.001%) and is a missense mutation (asparagine→glutamic acid). There is one additional synonymous variant associated with sPTB (rs17853270) in the 17th exon of *PPFIA1*. There are 156 variants linked to rs1601328, which compose a large and complex haplotype spanning approximately 129 kb. This variant also affects expression of two genes cortactin (*CTTN*) and *PPFIA1* in several tissues, including adipose, thyroid, and tibial nerve (GTEx v7)^73^.

Both *CTTN* and *PPFIA1* are involved in cell motility and cell adhesion. The PPFIA1 protein is a member of the LAR protein-tyrosine phosphatase-interacting protein (liprin) family and is involved in cell motility, extracellular matrix dynamics, and cell adhesion^74–77^. Cell adhesion and migration are also critical processes involved in placental development and implantation^78,79^ and other members of the liprin family have been linked to maternal-fetal signaling during placental development^80,81^. The CTTN protein is an actin-binding protein involved in cell migration and invasion^82–84^. *CTTN* was shown to be expressed in the decidual cells and spiral arterioles as well as localize to the trophoblast cells during early pregnancy—suggesting a role for *CTTN* in cytoskeletal remodeling of the maternal-fetal interface during early pregnancy^85,86^. While neither *CTTN* nor *PPFIA1* have been implicated in sPTB, they are both linked to cell adhesion and there is evidence that decreased adherence of maternal and fetal membrane layers is involved in parturition^87^. The accelerated evolution associated with variants in this region is also consistent with the rapid diversification of the placenta within eutherians ^88–90^. While the proteins CTTN and PPFIA1 are highly conserved across mammals, the accelerated evolution detected in association with rs1061328 may be linked to the functionally important and diverse splice variants of both proteins^74,91^. Accelerated evolution has previously been detected in the birth timing-associated genes *FSHR*^47^ and *PLA2G4C*^92^. It has been hypothesized that human and/or primate-specific adaptations, such as bipedalism, have resulted in the accelerated evolution of birth-timing phenotypes along these lineages^2,93,94^. Accelerated evolution has also been implicated in other complex disorders—especially those like schizophrenia^95,96^ and autism^97^ which affect the brain, another organ that is thought to have undergone adaptive evolution in the human lineage.

### Group D: Balanced Polymorphism/Balancing Selection

Variants in Group D generally had higher than expected values of beta score or an older allele age than expected, which is consistent with evolutionary signatures of balancing selection (Figure 2). Beta score is a measure of long-term balancing selection within the human lineage^33^. Allele age is measured as the time to most recent common ancestor (TMRCA) in thousands of generations since present based on 54 unrelated individuals, which was obtained from ARGweaver^34^. There are nine genomic regions in group D, of which three have a significantly (p<0.05) higher beta score than expected, three have a significantly older (p<0.05) than expected TMRCA, and three have older TMRCA but are not significant. The derived alleles in this group have an average derived allele frequency across all populations of 0.44 (Figure 3C).There are 292 variants in this group; nearly all of these variants are intronic (274/292; Figure 3A) and there is evidence of regulatory binding for 11 variants (regulome DB score of 1 or 2; Figure 3B). GTEx analysis supports the regulatory role of a number of variants—266 of 271 variants are an eQTL in at least one tissue (Figure 3D). For the genomic region associated with the lead variant rs10932774 there are 26 additional variants which are eQTLs for an average of 80 unique tissue/gene combinations. eQTL analysis detects at least one expression change in the uterus in all but five of these variants.

The variant rs10932774 is located within the 2^nd^ of nine introns in the gene *PNKD* (also known as myofibrillogenesis regulator 1), which is the causal gene in the neurological movement disorder paroxysmal nonkinesiogenic dyskinesia (PNKD)^98^. This variant has a significant value for both TMRCA and beta score. The G allele at this locus is associated with an increased risk of sPTB (OR: 1.11, adjusted p-value 8.85×10^−5^)^32^ and is the major allele in all the populations examined (Figure 4D). Supporting the interpretation of long-term balancing selection, this polymorphism is also present in the great apes. This variant is associated with the expression of five genes in 43 tissues for a total of 73 gene/tissue combinations in the GTEx database. Additionally, the *PNKD* gene is up-regulated in severely preeclamptic placentas^99^ and in PNKD patients pregnancy is associated with changes in the frequency or severity of PNKD attacks^100–102^. Expression changes in the Transmembrane BAX Inhibitor Motif-Containing Protein 1 gene (*TMBIM1)* and the Actin Related Protein 2/3 Complex Subunit 2 gene (*ARPC2*) are also associated with this variant. TMBIM1 is a cell death regulator^103^ with no known role in pregnancy. Methylation of *TMBIM1*, however, is altered in the offspring of mothers with Type 1 Diabetes^104^. The gene *ARPC2* is a subunit of the Arp2/3 complex which controls actin polymerization and is also highly conserved^105,106^. The complex is important for early embryo development and preimplantation in pigs and mice^107,108^. *ARPC2* expression has been identified in the BeWo trophoblastic cell line used to investigate placental function^109^ and is subject to RNA editing in placentas associated with intrauterine growth restriction/small for gestational age (SGA)^110^. Overall, genes associated with the variant rs10932774 (*PNKD, TMBIM1* and *ARPC2*) show long-term evolutionary conservation consistent with a signature of balancing selection and prior research suggests links to pregnancy through a variety of mechanisms. The identification of balancing selection acting on sPTB-associated genomic regions is consistent with the critical role of the immune system, which often experiences balancing selection^33,111,112^, in establishing and maintaining pregnancy^113^.

### The largest group of variants consists of a variety of evolutionary signatures

The final group, group E, contained the remaining genomic regions in clusters E1, E2, E3 and E4 and was associated with a broad range of evolutionary signatures (Figure 2). At least one variant in group E had a significant p-value for every evolutionary measure except for alignment block age. While this group does not reflect the action of a single evolutionary measure or force, over half of the lead variants (39/73) had a significant p-value (p<0.05) for either GERP or XP-EHH. This group also contained 23 of the 33 genomic regions with a z-score (|z|>1.5) for population specific iHS (Supp. Table 2). These genomic regions contained a total of 444 linked variants of which 242 are intronic variants, 178 are intergenic variants and the remaining 24 variants are upstream and downstream variants (Figure 4A). This group also contains 19 variants that are likely to affect binding (regulomeDB score of 1 or 2; Fig. 4b). The majority of the derived alleles in this group are minor in Europeans (313/444; Figure 4C). There are also 143 variants identified as eQTLs, including 16 expression changes for genes in the uterus (all associated with the variant rs12646130; Figure 4D). Interestingly, this group contained variants linked to the *EEFSEC, ADCY5*, and *WNT4* genes, which have been previously associated with gestational duration or preterm birth^32^.

The variant rs8126001 is located in the 5’ UTR of the opioid related nociception receptor 1 or nociception opioid receptor (*OPRL1* or *NOP-R*) gene which may be involved in myometrial contractions during delivery^114,115^. This variant has signatures of positive selection as detected by the integrated haplotype score (iHS) within the African population (Supp. Table 2). The T allele at this locus is protective for sPTB (effect: 0.896; adjusted p-value 4.04×10^−5^)^32^ and arose in the human lineage. The protective allele has a relatively low frequency in the African population and is located in a region with low haplotype diversity (Figure 4E). This locus is also associated with expression of *OPRL1* in tissues like the brain, aorta and esophagus (Figure 4E). The gene *OPRL1* is a receptor for the endogenous peptide nociceptin (N/OFQ) which is derived from prenocicpetin (PNOC). There is evidence that nociception and its receptor may play a role in pregnancy. N/OFQ and PNOC were detected in rat and human pregnant myometrial tissues^32^ and *OPRL1* was detected in rat myometrium.^114,116^ Additionally, *PNOC* mRNA levels are significantly higher in preterm uterine samples in humans and can elicit myometrial relaxation *in vitro*^115,116^. It is therefore likely that nociceptin and *OPRL1* are involved in the perception of pain during delivery and the initiation of delivery. While a single mode of evolution does not characterize group E, the high frequency of genomic regions with significant XP-EHH or population specific iHS values (40/73 genomic regions) suggests that population-specific evolutionary forces may be at play in this group. The sPTB GWAS and population-specific evolutionary measures were conducted in women of European ancestry but we know that sPTB risk varies with genomic background^117,118^. Therefore, this group also suggests that individual populations experience a different mosaic of evolutionary forces on pregnancy phenotypes.

## CONCLUSIONS

In this study, we developed an approach to test for signatures of diverse evolutionary forces and applied it to sPTB-associated genomic regions. This approach explicitly accounts for MAF and LD in trait-associated genomic regions. We find evolutionary conservation, excess population differentiation, accelerated evolution, and balanced polymorphisms in 120 of the 215 sPTB-associated genomic regions. Annotation of these regions using existing databases and literature suggest plausible functional links to pregnancy phenotypes, bolstering our confidence that these regions contribute to sPTB risk. These results suggest that no single evolutionary force is responsible for shaping the genetic architecture of sPTB; rather, sPTB has been influenced by a diverse mosaic of evolutionary forces. We hypothesize that the same is likely to be true of other complex human traits and disorders; future investigations that test for signatures of multiple evolutionary forces, such as ours, promise to elucidate the degree to which the landscape of evolutionary forces varies across disorders.

## METHODS

### Deriving independent genomic regions associated with sPTB from GWAS summary statistics

To evaluate evolutionary history of sPTB on distinct regions of the human genome, we identified genomic regions from the GWAS summary statistics. Using PLINK1.9b (pngu.mgh.harvard.edu/purcell/plink/)^102^, the top 10,000 variants associated with sPTB from Zhang et. al.^31^ were clumped based on LD using default settings except requiring a p-value ≤ 10E-4 for lead variants and variants in LD with lead variants. We used this liberal p-value threshold to increase the number of sPTB-associated variants evaluated. Although this will increase the number of false positive variants associated with sPTB, we anticipate that these false positive variants will not have statistically significant evolutionary signals using our approach to detect evolutionary forces. This is because the majority of the genome is neutrally evolving and our approach aims to detect deviation from this genomic background. Additionally, it is possible that the lead variant (variant with the lowest p-value) could tag the true variant associated with sPTB within an LD block. Therefore, we defined an independent sPTB-associated genomic region to include the lead and LD (r^2^>0.9, p-value <= 10E-4) sPTB variants. This resulted in 215 independent lead variants within an sPTB-associated genomic region.

### Creating matched control regions for each sPTB-associated genomic regions

We detected evolutionary signatures at genomic regions associated with sPTB by comparing them to matched control sets. Since evolutionary measures are influenced by LD and allele frequencies and these also influence power in GWAS, we generated control regions matched for these attributes for observed sPTB-associated genomic regions. First, for each lead variant we identified 5,000 control variants matched on minor allele frequency (+/-5%), LD (r^2^>0.9, +/-10% number of LD buddies), gene density (+/- 500%) and distance to nearest gene (+/-500%) using SNPSNAP^119^, which derives controls variants from a quality controlled phase 3 100 Genomes (1KG) data, with default settings for all other parameters and the hg19/GRCh37 genome assembly. For each control variant, we randomly selected an equal number of variants in LD (r^2^>0.9) as sPTB-associated variants in LD with the corresponding lead variant. If no matching control variant existed, we relaxed the LD required to r^2^=0.6. If still no match was found, we treated this as a missing value. For all LD calculations, control variants and downstream evolutionary measure analyses, the European super-population from phase 3 1KG^120^ was used after removing duplicate variants.

### Evolutionary measures

To characterize the evolutionary dynamics at each sPTB-associated region, we evaluated diverse evolutionary measures for diverse modes of selection and allele history across each sPTB-associated genomic region. Evolutionary measures were either calculated or pre-calculated values were downloaded for all control and sPTB-associated variants. Pairwise Weir and Cockerham’s F_ST_ values between European, East Asian, and African super populations from 1KG were calculated using VCFTools (v0.1.14)^121^. Evolutionary measures of positive selection, iHS, XP-EHH, and iES, were calculated from the 1KG data using rehh 2.0^122^. Beta score, a measure of balancing selection, was calculated using BetaScan software^33^. Alignment block age was calculated using 100-way multiple sequence alignment^123^ to measure the age of alignment blocks defined by the oldest most recent common ancestor. The remaining measures were downloaded from publicly available sources: phyloP and phastCons 100 way alignment from UCSC genome browser^124^; LINSIGHT^35^; and allele age (time to most common recent ancestor from ARGWEAVER)^34^. Due to missing values, the exact number of control regions varied by sPTB-associated region and evolutionary measure. We first marked any control set that did not match at least 90% of the required variants for a given sPTB-associated region, then any sPTB-associated region with ≥ 60% marked control regions were removed for that specific evolutionary measure. iHS was not included in Figure 2 because of large amounts of missing data for up to 50% of genomic regions evaluated.

### Detecting significant differences in evolutionary measures by comparing to control distributions

For each sPTB-associated genomic region for a specific evolutionary measure, we took the median value of the evolutionary measure across all its variants and compared it to the distribution of median values from the corresponding control regions. Statistical significance for each sPTB-associated region was evaluated by comparing the median value of the evolutionary measure to the distribution of the median value of the control regions. To obtain the p-value, we calculated the number of control regions with a median value that are equal to or greater the median value for the PTB region. Since allele age (TMRCA from ARGweaver), PhyloP, and alignment block age are bi-directional measures, we calculated two-tailed p-values; all other evolutionary measures used one-tailed p-values. To compare evolutionary measures whose scales differ substantially, we calculated a z-score for each region per measure. These z-scores were hierarchically clustered across all regions and measures. Clusters were defined by a branch length cutoff of seven. These clusters were then grouped and annotated by the dominant evolutionary measure through manual inspection to highlight the main evolutionary trend(s).

### Annotation of variants in sPTB-associated regions

To understand functional differences between groups and genomic regions we collected annotations for variants in sPTB-associated regions from publicly available databases. Evidence for regulatory function for individual variants was obtained from RegulomeDB v1.1 (accessed 1/11/19)^125^. From this we extracted the following information: total promotor histone marks, total enhancer histone marks, total DNase 1 sensitivity, total predicted proteins bound, total predicted motifs changed, and regulomeDB score. Variants were identified as expression quantitative trait loci (eQTLs) using the Genotype-Tissue Expression (GTEx) project data (dbGaP Accession phs000424.v7.p2 accessed 1/15/19). Variants were mapped to GTEx annotations based on rs number and then the GTEx annotations were used to obtain eQTL information. For each locus, we obtained the tissues in which the locus was an eQTL, the genes for which the locus affected expression (in any tissue), and the total number times the locus was identified as an eQTL. Functional variant effects were annotated with the Ensembl Variant Effect Predictor (VEP; accessed 1/17/19) based on rs number^126^. Population-based allele frequencies were obtained from the 1KG phase3 data for the African (excluding related African individuals; Supplementary Table 3), East Asian, and European populations^120^.

To infer the history of the alleles at each locus across mammals, we created a mammalian alignment at each locus and inferred the ancestral states. That mammalian alignment was built using data from the sPTB GWAS^32^ (risk variant identification), the UCSC Table Browser^123^ (30 way mammalian alignment), the 1KG phase 3^120^ data (human polymorphism data) and the Great Ape Genome project (great ape polymorphisms)^127^—which reference different builds of the human genome. To access data constructed across multiple builds of the human genome, we used Ensembl biomart release 97^128^ and the biomaRt R package^129,130^ to obtain the position of variants in hg38, hg19, and hg18 based on RefSNP (rs) number^131^. Alignments with more than one gap position were discarded due to uncertainty in the alignment. All variant data were checked to ensure that each dataset reported polymorphisms in reference to the same strand. Parsimony reconstruction was conducted along a phylogenetic tree generated from the TimeTree database^132^. Ancestral state reconstruction for each allele was conducted in R using parsimony estimation in the phangorn package^133^. Five character-states were used in the ancestral state reconstruction: one for each base and a fifth for gap. Haplotype blocks containing the variant of interest were identified using Plink (v1.9b_5.2) to create blocks from the 1KG phase3 data. Binary haplotypes were then generated for each of the three populations using the IMPUTE function of vcftools (v0.1.15.) Median joining networks^134^ were created using PopART^135^.

## URLs

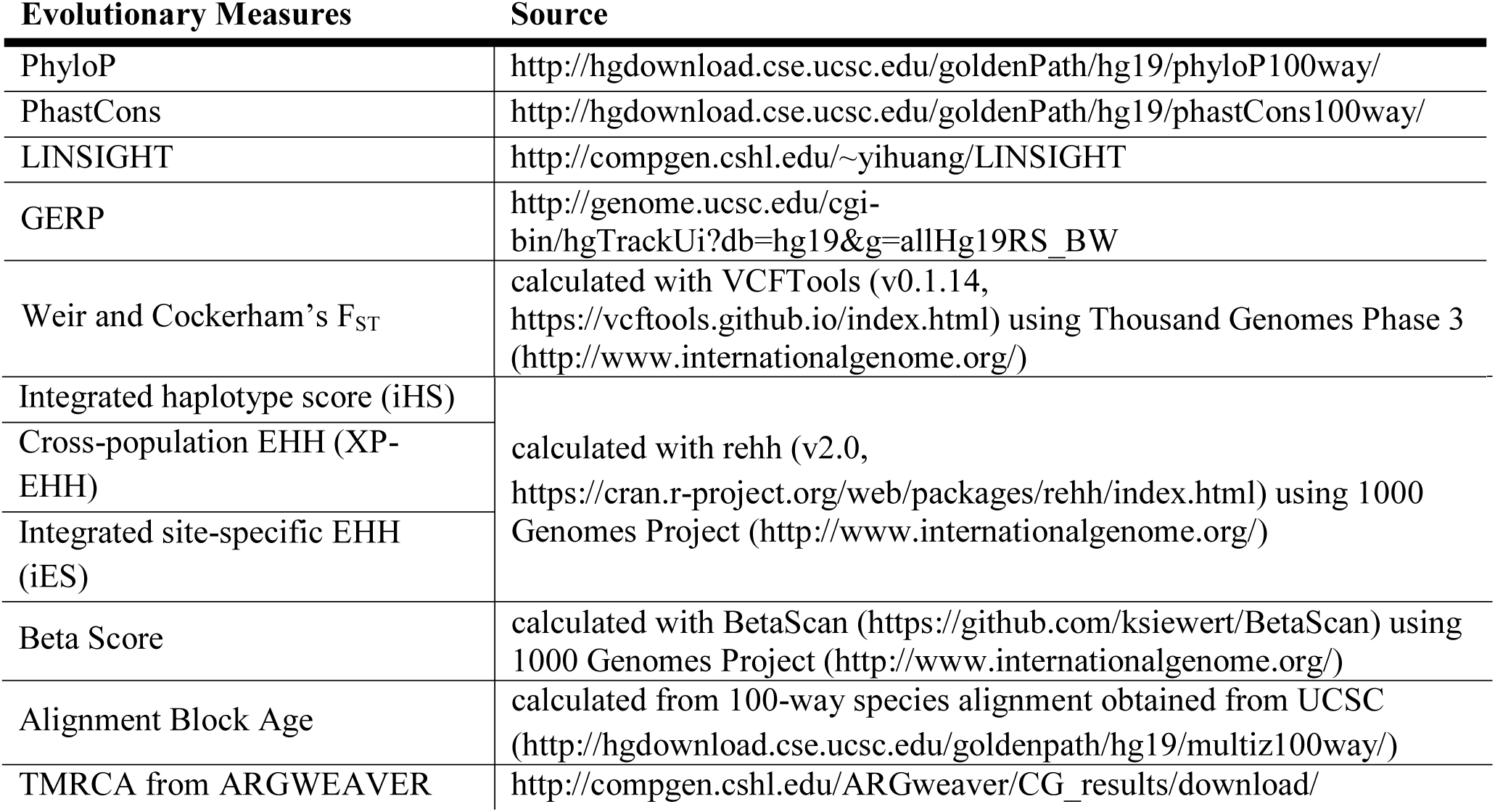

## Data Availability

All the data used in this study were obtained from the public domain (see the URLs above) or deposited in a figshare repository to be made public upon publication.

## Code Availability

All scripts used to measure evolutionary signatures and generate figures are publicly accessible in a figshare repository to be made public upon publication.

## Acknowledgements

This work was supported by the National Institutes of Health (grant R35GM127087 to JAC), the Burroughs Wellcome Fund Preterm Birth Initiative (to JAC and AR), and by the March of Dimes through the March of Dimes Prematurity Research Center Ohio Collaborative (to LJM, GZ, PA, JAC, and AR). AA was also supported by NIGMS of the National Institutes of Health under award number T32GM007347. This work was conducted in part using the resources of the Advanced Computing Center for Research and Education at Vanderbilt University. The content is solely the responsibility of the authors and does not necessarily represent the official views of the National Institutes of Health, the March of Dimes, or the Burroughs Wellcome Fund.

## Author Contributions

A.A., A.L.L., P.A., J.A.C, A.R. conceived and designed the study. A.A. and A.L.L performed all statistical analyses, functional annotations, and wrote the manuscript under guidance from P.A., J.A.C, A.R. G.Z. and L.M provided sPTB-associated genomic regions, guidance, and feedback on the manuscript. Y.P. calculated the beta score measure for balancing selection using BetaScan^33^. S.F. calculated the alignment block age. All authors reviewed and approved the final manuscript.

## References

1. Eidem, H. R., McGary, K. L., Capra, J. A., Abbot, P. & Rokas, A. The transformative potential of an integrative approach to pregnancy. Placenta 57, 204–215 (2017).

2. Abbot, P. & Rokas, A. Mammalian pregnancy. Current Biology (2017). doi:10.1016/j.cub.2016.10.046

3. Moon, J. M., Capra, J. A., Abbot, P. & Rokas, A. Immune Regulation in Eutherian Pregnancy: Live Birth Coevolved with Novel Immune Genes and Gene Regulation. BioEssays (2019). doi:10.1002/bies.201900072

4. Rosenberg, K. & Trevathan, W. Bipedalism and human birth: The obstetrical dilemma revisited. Evol. Anthropol. Issues, News, Rev. 4, 161–168 (1995).

5. Rosenberg, K. R. The evolution of modern human childbirth. Am. J. Phys. Anthropol. (1992). doi:10.1002/ajpa.1330350605

6. Washburn, S. L. Tools and human evolution. Sci Am 203, 63–75 (1960).

7. Krogman, W. M. The scars of human evolution. Sci. Am. 185, 54–57 (1951).

8. Dunsworth, H. M., Warrener, A. G., Deacon, T., Ellison, P. T. & Pontzer, H. Metabolic hypothesis for human altriciality. Proc Natl Acad Sci U S A 109, 15212–15216 (2012).

9. Romero, R., Dey, S. K. & Fisher, S. J. Preterm labor: One syndrome, many causes. Science (2014). doi:10.1126/science.1251816

10. Martin, J. A., Hamilton, B. E. & Osterman, M. J. K. Births in the United States, 2016. NCHS Data Brief (2017).

11. Blencowe, H. et al. National, regional, and worldwide estimates of preterm birth rates in the year 2010 with time trends since 1990 for selected countries: A systematic analysis and implications. Lancet (2012). doi:10.1016/S0140-6736(12)60820-4

12. Goldenberg, R. L., Culhane, J. F., Iams, J. D. & Romero, R. Epidemiology and causes of preterm birth. The Lancet (2008). doi:10.1016/S0140-6736(08)60074-4

13. Esplin, M. S. Overview of spontaneous preterm birth: A complex and multifactorial phenotype. In Clinical Obstetrics and Gynecology (2014). doi:10.1097/GRF.0000000000000037

14. Chang, H. H. et al. Preventing preterm births: Analysis of trends and potential reductions with interventions in 39 countries with very high human development index. Lancet (2013). doi:10.1016/S0140-6736(12)61856-X

15. Bezold, K. Y., Karjalainen, M. K., Hallman, M., Teramo, K. & Muglia, L. J. The genomics of preterm birth: From animal models to human studies. Genome Medicine (2013). doi:10.1186/gm438

16. Moutquin, J. M. Classification and heterogeneity of preterm birth. in BJOG: An International Journal of Obstetrics and Gynaecology (2003). doi:10.1016/S1470-0328(03)00021-1

17. Barros, F. C. et al. The Distribution of Clinical Phenotypes of Preterm Birth Syndrome. JAMA Pediatr. (2015). doi:10.1001/jamapediatrics.2014.3040

18. Ananth, C. V. & Vintzileos, A. M. Epidemiology of preterm birth and its clinical subtypes. Journal of Maternal-Fetal and Neonatal Medicine (2006). doi:10.1080/14767050600965882

19. Henderson, J. J., McWilliam, O. A., Newnham, J. P. & Pennell, C. E. Preterm birth aetiology 2004-2008. Maternal factors associated with three phenotypes: Spontaneous preterm labour, preterm pre-labour rupture of membranes and medically indicated preterm birth. J. Matern. Neonatal Med. (2012). doi:10.3109/14767058.2011.597899

20. Boyd, H. A. et al. Maternal Contributions to Preterm Delivery. Am. J. Epidemiol. (2009). doi:10.1093/aje/kwp324

21. Kistka, Z. A. F. et al. Heritability of parturition timing: an extended twin design analysis. Am. J. Obstet. Gynecol. (2008). doi:10.1016/j.ajog.2007.12.014

22. Plunkett, J. et al. Mother’s genome or maternally-inherited genes acting in the fetus influence gestational age in familial preterm birth. Hum. Hered. (2009). doi:10.1159/000224641

23. York, T. P. et al. Fetal and maternal genes’ influence on gestational age in a quantitative genetic analysis of 244,000 swedish births. Am. J. Epidemiol. (2013). doi:10.1093/aje/kwt005

24. Clausson, B., Lichtenstein, P. & Cnattingius, S. Genetic influence on birthweight and gestational length determined by studies in offspring of twins. BJOG An Int. J. Obstet. Gynaecol. (2000). doi:10.1111/j.1471-0528.2000.tb13234.x

25. Kjeldbjerg, A. L., Villesen, P., Aagaard, L. & Pedersen, F. S. Gene conversion and purifying selection of a placenta-specific ERV-V envelope gene during simian evolution. BMC Evol Biol 8, 266 (2008).

26. Hiby, S. E. et al. Maternal KIR in Combination with Paternal HLA-C2 Regulate Human Birth Weight. J. Immunol. (2014). doi:10.4049/jimmunol.1400577

27. Guinan, K. J. et al. Signatures of natural selection and coevolution between killer cell immunoglobulin-like receptors (KIR) and HLA class i genes. Genes Immun. (2010). doi:10.1038/gene.2010.9

28. Phillips, J. B., Abbot, P. & Rokas, A. Is preterm birth a human-specific syndrome? Evol. Med. Public Heal. (2015). doi:10.1093/emph/eov010

29. Chen, C. et al. The human progesterone receptor shows evidence of adaptive evolution associated with its ability to act as a transcription factor. Mol. Phylogenet. Evol. (2008). doi:10.1016/j.ympev.2007.12.026

30. Li, J. et al. Natural Selection Has Differentiated the Progesterone Receptor among Human Populations. Am J Hum Genet 103, 45–57 (2018).

31. Newnham, J. P. et al. Strategies to prevent preterm birth. Frontiers in Immunology (2014). doi:10.3389/fimmu.2014.00584

32. Zhang, G., Jacobsson, B. & Muglia, L. J. Genetic Associations with Spontaneous Preterm Birth. N. Engl. J. Med. 377, 2401–2402 (2017).

33. Siewert, K. M. & Voight, B. F. Detecting Long-Term Balancing Selection Using Allele Frequency Correlation. Mol Biol Evol 34, 2996–3005 (2017).

34. Rasmussen, M. D., Hubisz, M. J., Gronau, I. & Siepel, A. Genome-Wide Inference of Ancestral Recombination Graphs. PLoS Genet. (2014). doi:10.1371/journal.pgen.1004342

35. Huang, Y. F., Gulko, B. & Siepel, A. Fast, scalable prediction of deleterious noncoding variants from functional and population genomic data. Nat. Genet. (2017). doi:10.1038/ng.3810

36. Akey, J. M. Constructing genomic maps of positive selection in humans: Where do we go from here? Genome Research (2009). doi:10.1101/gr.086652.108

37. Pybus, M. et al. 1000 Genomes Selection Browser 1.0: A genome browser dedicated to signatures of natural selection in modern humans. Nucleic Acids Res. (2014). doi:10.1093/nar/gkt1188

38. Vitti, J. J., Grossman, S. R. & Sabeti, P. C. Detecting Natural Selection in Genomic Data. Annu. Rev. Genet. (2013). doi:10.1146/annurev-genet-111212-133526

39. Stern, A. J. & Nielsen, R. Detecting Natural Selection. Handb. Stat. Genomics 4e 2V SET 340–397 (2019).

40. Booker, T. R., Jackson, B. C. & Keightley, P. D. Detecting positive selection in the genome. BMC Biology (2017). doi:10.1186/s12915-017-0434-y

41. Visscher, P. M. et al. 10 Years of GWAS Discovery: Biology, Function, and Translation. American Journal of Human Genetics (2017). doi:10.1016/j.ajhg.2017.06.005

42. Guo, J. et al. Global genetic differentiation of complex traits shaped by natural selection in humans. Nat. Commun. (2018). doi:10.1038/s41467-018-04191-y

43. Byars, S. G. et al. Genetic loci associated with coronary artery disease harbor evidence of selection and antagonistic pleiotropy. PLoS Genet. (2017). doi:10.1371/journal.pgen.1006328

44. O’Connor, L. J. et al. Extreme Polygenicity of Complex Traits Is Explained by Negative Selection. Am. J. Hum. Genet. (2019). doi:10.1016/j.ajhg.2019.07.003

45. Zeng, J. et al. Signatures of negative selection in the genetic architecture of human complex traits. Nat. Genet. (2018). doi:10.1038/s41588-018-0101-4

46. Guo, J., Yang, J. & Visscher, P. M. Leveraging GWAS for complex traits to detect signatures of natural selection in humans. Current Opinion in Genetics and Development (2018). doi:10.1016/j.gde.2018.05.012

47. Plunkett, J. et al. An evolutionary genomic approach to identify genes involved in human birth timing. PLoS Genet 7, e1001365 (2011).

48. Guinan, K. J. et al. Signatures of natural selection and coevolution between killer cell immunoglobulin-like receptors (KIR) and HLA class I genes. Genes Immun 11, 467–478 (2010).

49. Gu, T. P. et al. The role of Tet3 DNA dioxygenase in epigenetic reprogramming by oocytes. Nature 477, 606–610 (2011).

50. Tsukada, Y., Akiyama, T. & Nakayama, K. I. Maternal TET3 is dispensable for embryonic development but is required for neonatal growth. Sci Rep 5, 15876 (2015).

51. Rakoczy, J. et al. Dynamic expression of TET1, TET2, and TET3 dioxygenases in mouse and human placentas throughout gestation. Placenta 59, 46–56 (2017).

52. Sober, S. et al. Extensive shift in placental transcriptome profile in preeclampsia and placental origin of adverse pregnancy outcomes. Sci Rep 5, 13336 (2015).

53. Fitzgerald, E., Boardman, J. P. & Drake, A. J. Preterm Birth and the Risk of Neurodevelopmental Disorders – Is There a Role for Epigenetic Dysregulation? Curr Genomics 19, 507–521 (2018).

54. Zelko, I. N., Zhu, J. & Roman, J. Maternal undernutrition during pregnancy alters the epigenetic landscape and the expression of endothelial function genes in male progeny. Nutr Res 61, 53–63 (2019).

55. Akahori, H., Guindon, S., Yoshizaki, S. & Muto, Y. Molecular evolution of the TET gene family in mammals. Int. J. Mol. Sci. (2015). doi:10.3390/ijms161226110

56. Gazal, S. et al. Linkage disequilibrium-dependent architecture of human complex traits shows action of negative selection. Nat. Genet. (2017). doi:10.1038/ng.3954

57. Kistka, Z. A. et al. Racial disparity in the frequency of recurrence of preterm birth. Am J Obs. Gynecol 196, 131 e1–6 (2007).

58. Muglia, L. J. & Katz, M. The enigma of spontaneous preterm birth. N Engl J Med 362, 529–535 (2010).

59. Jung, K. H. et al. Associations of vitamin d binding protein gene polymorphisms with the development of peripheral arthritis and uveitis in ankylosing spondylitis. J Rheumatol 38, 2224–2229 (2011).

60. Muindi, J. R. et al. Serum vitamin D metabolites in colorectal cancer patients receiving cholecalciferol supplementation: correlation with polymorphisms in the vitamin D genes. Horm Cancer 4, 242–250 (2013).

61. Wei, S. Q., Qi, H. P., Luo, Z. C. & Fraser, W. D. Maternal vitamin D status and adverse pregnancy outcomes: a systematic review and meta-analysis. J Matern Fetal Neonatal Med 26, 889–899 (2013).

62. Bodnar, L. M., Platt, R. W. & Simhan, H. N. Early-pregnancy vitamin D deficiency and risk of preterm birth subtypes. Obs. Gynecol 125, 439–447 (2015).

63. Qin, L. L., Lu, F. G., Yang, S. H., Xu, H. L. & Luo, B. A. Does Maternal Vitamin D Deficiency Increase the Risk of Preterm Birth: A Meta-Analysis of Observational Studies. Nutrients 8, (2016).

64. Zhou, S. S., Tao, Y. H., Huang, K., Zhu, B. B. & Tao, F. B. Vitamin D and risk of preterm birth: Up-to-date meta-analysis of randomized controlled trials and observational studies. J Obs. Gynaecol Res 43, 247–256 (2017).

65. Liong, S., Di Quinzio, M. K., Fleming, G., Permezel, M. & Georgiou, H. M. Is vitamin D binding protein a novel predictor of labour? PLoS One 8, e76490 (2013).

66. D’Silva, A. M., Hyett, J. A. & Coorssen, J. R. Proteomic analysis of first trimester maternal serum to identify candidate biomarkers potentially predictive of spontaneous preterm birth. J. Proteomics (2018). doi:10.1016/j.jprot.2018.02.002

67. Bodnar, L. M. & Simhan, H. N. Vitamin D may be a link to black-white disparities in adverse birth outcomes. Obs. Gynecol Surv 65, 273–284 (2010).

68. Burris, H. H. et al. Plasma 25-hydroxyvitamin D during pregnancy and small-for-gestational age in black and white infants. Ann Epidemiol 22, 581–586 (2012).

69. Reeves, I. V. et al. Vitamin D deficiency in pregnant women of ethnic minority: A potential contributor to preeclampsia. J. Perinatol. (2014). doi:10.1038/jp.2014.91

70. Ramagopalan, S. V. et al. A ChIP-seq defined genome-wide map of vitamin D receptor binding: Associations with disease and evolution. Genome Res. (2010). doi:10.1101/gr.107920.110

71. Jablonski, N. G. & Chaplin, G. The roles of vitamin D and cutaneous vitamin D production in human evolution and health. International Journal of Paleopathology (2018). doi:10.1016/j.ijpp.2018.01.005

72. Hollis, B. W. & Wagner, C. L. New insights into the vitamin D requirements during pregnancy. Bone Res 5, 17030 (2017).

73. Carithers, L. J. et al. A Novel Approach to High-Quality Postmortem Tissue Procurement: The GTEx Project. Biopreserv. Biobank. (2015). doi:10.1089/bio.2015.0032

74. Zurner, M. & Schoch, S. The mouse and human Liprin-alpha family of scaffolding proteins: Genomic organization, expression profiling and regulation by alternative splicing. Genomics 93, 243–253 (2009).

75. Asperti, C., Pettinato, E. & de Curtis, I. Liprin-alpha1 affects the distribution of low-affinity beta1 integrins and stabilizes their permanence at the cell surface. Exp Cell Res 316, 915–926 (2010).

76. Astro, V., Asperti, C., Cangi, M. G., Doglioni, C. & de Curtis, I. Liprin-alpha1 regulates breast cancer cell invasion by affecting cell motility, invadopodia and extracellular matrix degradation. Oncogene 30, 1841–1849 (2011).

77. de Curtis, I. Function of liprins in cell motility. Exp Cell Res 317, 1–8 (2011).

78. Yang, J. T., Rayburn, H. & Hynes, R. O. Cell adhesion events mediated by alpha 4 integrins are essential in placental and cardiac development. Development 121, 549–560 (1995).

79. Burrows, T. D., King, A. & Loke, Y. W. Trophoblast migration during human placental implantation. Hum Reprod Updat. 2, 307–321 (1996).

80. Mincheva-Nilsson, L. & Baranov, V. The role of placental exosomes in reproduction. Am J Reprod Immunol 63, 520–533 (2010).

81. Paidas, M. J. et al. A genomic and proteomic investigation of the impact of preimplantation factor on human decidual cells. Am J Obs. Gynecol 202, 459 e1–8 (2010).

82. Weed, S. A. & Parsons, J. T. Cortactin: coupling membrane dynamics to cortical actin assembly. Oncogene 20, 6418–6434 (2001).

83. van Rossum, A. G., Moolenaar, W. H. & Schuuring, E. Cortactin affects cell migration by regulating intercellular adhesion and cell spreading. Exp Cell Res 312, 1658–1670 (2006).

84. Clark, E. S., Whigham, A. S., Yarbrough, W. G. & Weaver, A. M. Cortactin is an essential regulator of matrix metalloproteinase secretion and extracellular matrix degradation in invadopodia. Cancer Res 67, 4227–4235 (2007).

85. Paule, S. G., Airey, L. M., Li, Y., Stephens, A. N. & Nie, G. Proteomic approach identifies alterations in cytoskeletal remodelling proteins during decidualization of human endometrial stromal cells. J Proteome Res 9, 5739–5747 (2010).

86. Paule, S., Li, Y. & Nie, G. Cytoskeletal remodelling proteins identified in fetal-maternal interface in pregnant women and rhesus monkeys. J Mol Histol 42, 161–166 (2011).

87. Strohl, A. et al. Decreased adherence and spontaneous separation of fetal membrane layers--amnion and choriodecidua--a possible part of the normal weakening process. Placenta 31, 18–24 (2010).

88. Ford, S. P. Embryonic and fetal development in different genotypes in pigs. J. Reprod. Fertil. Suppl. (1997).

89. Mossman, H. W. Comparative morphogenesis of the fetal membranes and accessory uterine structures. Placenta (1991). doi:10.1016/0143-4004(91)90504-9

90. Chuong, E. B., Hannibal, R. L., Green, S. L. & Baker, J. C. Evolutionary perspectives into placental biology and disease. Applied and Translational Genomics (2013). doi:10.1016/j.atg.2013.07.001

91. van Rossum, A. G. S. H., Schuuring-Scholtes, E., van Buuren-van Seggelen, V., Kluin, P. M. & Schuuring, E. Comparative genome analysis of cortactin and HS1: The significance of the F-actin binding repeat domain. BMC Genomics (2005). doi:10.1186/1471-2164-6-15

92. Plunkett, J. et al. Primate-specific evolution of noncoding element insertion into PLA2G4C and human preterm birth. BMC Med. Genomics (2010). doi:10.1186/1755-8794-3-62

93. Rosenberg, K. & Trevathan W. Birth, obstetrics and human evolution. BJOG: An International Journal of Obstetrics and Gynaecology (2002). doi:10.1046/j.1471-0528.2002.00010.x

94. Weaver, T. D. & Hublin, J. J. Neandertal birth canal shape and the evolution of human childbirth. Proc. Natl. Acad. Sci. U. S. A. (2009). doi:10.1073/pnas.0812554106

95. Xu, K., Schadt, E. E., Pollard, K. S., Roussos, P. & Dudley, J. T. Genomic and network patterns of schizophrenia genetic variation in human evolutionary accelerated regions. Mol. Biol. Evol. (2015). doi:10.1093/molbev/msv031

96. Srinivasan, S. et al. Genetic Markers of Human Evolution Are Enriched in Schizophrenia. Biol. Psychiatry (2016). doi:10.1016/j.biopsych.2015.10.009

97. Polimanti, R. & Gelernter, J. Widespread signatures of positive selection in common risk alleles associated to autism spectrum disorder. PLoS Genet. (2017). doi:10.1371/journal.pgen.1006618

98. Rainier, S. et al. Myofibrillogenesis regulator 1 gene mutations cause paroxysmal dystonic choreoathetosis. Arch. Neurol. (2004). doi:10.1001/archneur.61.7.1025

99. Sitras, V. et al. Differential Placental Gene Expression in Severe Preeclampsia. Placenta (2009). doi:10.1016/j.placenta.2009.01.012

100. Stefano, E. et al. Clinical characteristics of paroxysmal nonkinesigenic dyskinesia in Serbian family with Myofibrillogenesis regulator 1 gene mutation. Mov. Disord. (2006). doi:10.1002/mds.21095

101. Ghezzi, D. et al. A family with paroxysmal nonkinesigenic dyskinesias (PNKD): Evidence of mitochondrial dysfunction. Eur. J. Paediatr. Neurol. (2015). doi:10.1016/j.ejpn.2014.10.003

102. Friedman, A. et al. Paroxysmal non-kinesigenic dyskinesia caused by the mutation of MR-1 in a large polish kindred. Eur. Neurol. (2008). doi:10.1159/000165348

103. Xu, Q. & Reed, J. C. Bax inhibitor-1, a mammalian apoptosis suppressor identified by functional screening in yeast. Mol. Cell (1998). doi:10.1016/S1097-2765(00)80034-9

104. Gautier, J. F. et al. Kidney dysfunction in adult offspring exposed in utero to type 1 diabetes is associated with alterations in genome-wide DNA methylation. PLoS One (2015). doi:10.1371/journal.pone.0134654

105. Welch, M. D., Iwamatsu, A. & Mitchison, T. J. Actin polymerization is induced by Arp2/3 protein complex at the surface of Listeria monocytogenes. Nature (1997). doi:10.1038/385265a0

106. Machesky, L. M., Atkinson, S. J., Ampe, C., Vandekerckhove, J. & Pollard, T. D. Purification of a cortical complex containing two unconventional actins from Acanthamoeba by affinity chromatography on profilin-agarose. J. Cell Biol. (1994). doi:10.1083/jcb.127.1.107

107. Sun, S.-C. et al. Actin nucleator Arp2/3 complex is essential for mouse preimplantation embryo development. Reprod. Fertil. Dev. (2013). doi:10.1071/rd12011

108. Li, Y. H. et al. Inhibition of the Arp2/3 complex impairs early embryo development of porcine parthenotes. Animal Cells Syst. (Seoul). (2016). doi:10.1080/19768354.2016.1228545

109. Szklanna, P. B. et al. Comparative proteomic analysis of trophoblast cell models reveals their differential phenotypes, potential uses, and limitations. Proteomics (2017). doi:10.1002/pmic.201700037

110. Majewska, M. et al. Placenta transcriptome profiling in intrauterine growth restriction (IUGR). Int. J. Mol. Sci. (2019). doi:10.3390/ijms20061510

111. Ferrer-Admetlla, A. et al. Balancing Selection Is the Main Force Shaping the Evolution of Innate Immunity Genes. J. Immunol. (2008). doi:10.4049/jimmunol.181.2.1315

112. Andrés, A. M. et al. Targets of balancing selection in the human genome. Mol. Biol. Evol. (2009). doi:10.1093/molbev/msp190

113. Mor, G. & Cardenas, I. The Immune System in Pregnancy: A Unique Complexity. American Journal of Reproductive Immunology (2010). doi:10.1111/j.1600-0897.2010.00836.x

114. Klukovits, A. et al. Nociceptin Inhibits Uterine Contractions in Term-Pregnant Rats by Signaling Through Multiple Pathways1. Biol. Reprod. (2010). doi:10.1095/biolreprod.109.082222

115. Gáspár, R., Deák, B. H., Klukovits, A., Ducza, E. & Tekes, K. Effects of Nociceptin and Nocistatin on Uterine Contraction. in Vitamins and Hormones (2015). doi:10.1016/bs.vh.2014.10.004

116. Bh, D. Uterus-Relaxing Effects of Nociceptin and Nocistatin: Studies on Preterm and Term-Pregnant Human Myometrium In vitro. Reprod. Syst. Sex. Disord. (2013). doi:10.4172/2161-038x.1000117

117. Manuck, T. A. et al. Admixture mapping to identify spontaneous preterm birth susceptibility loci in African Americans. Obstet. Gynecol. (2011). doi:10.1097/AOG.0b013e318214e67f

118. York, T. P., Eaves, L. J., Neale, M. C. & Strauss, J. F. The contribution of genetic and environmental factors to the duration of pregnancy. American Journal of Obstetrics and Gynecology (2014). doi:10.1016/j.ajog.2013.10.001

119. Pers, T. H., Timshel, P. & Hirschhorn, J. N. SNPsnap: A Web-based tool for identification and annotation of matched SNPs. Bioinformatics (2015). doi:10.1093/bioinformatics/btu655

120. Genomes Project, C. et al. A global reference for human genetic variation. Nature 526, 68–74 (2015).

121. Danecek, P. et al. The variant call format and VCFtools. Bioinformatics (2011). doi:10.1093/bioinformatics/btr330

122. Gautier, M., Klassmann, A. & Vitalis, R. rehh 2.0: a reimplementation of the R package rehh to detect positive selection from haplotype structure. Mol Ecol Resour 17, 78–90 (2017).

123. Karolchik, D. et al. The UCSC Table Browser data retrieval tool. Nucleic Acids Res. 32, D493–D496 (2004).

124. Kent, W. J. et al. The human genome browser at UCSC. Genome Res 12, 996–1006 (2002).

125. Boyle, A. P. et al. Annotation of functional variation in personal genomes using RegulomeDB. Genome Res 22, 1790–1797 (2012).

126. McLaren, W. et al. The Ensembl Variant Effect Predictor. Genome Biol 17, 122 (2016).

127. Prado-Martinez, J. et al. Great ape genetic diversity and population history. Nature 499, 471–475 (2013).

128. Zerbino, D. R. et al. Ensembl 2018. Nucleic Acids Res 46, D754–D761 (2018).

129. Durinck, S. et al. BioMart and Bioconductor: a powerful link between biological databases and microarray data analysis. Bioinformatics 21, 3439–3440 (2005).

130. Durinck, S., Spellman, P. T., Birney, E. & Huber, W. Mapping identifiers for the integration of genomic datasets with the R/Bioconductor package biomaRt. Nat Protoc 4, 1184–1191 (2009).

131. Lander, E. S. et al. Initial sequencing and analysis of the human genome. Nature 409, 860–921 (2001).

132. Kumar, S., Stecher, G., Suleski, M. & Hedges, S. B. TimeTree: A Resource for Timelines, Timetrees, and Divergence Times. Mol Biol Evol 34, 1812–1819 (2017).

133. Schliep, K. P. phangorn: phylogenetic analysis in R. Bioinformatics 27, 592–593 (2011).

134. Bandelt, H. J., Forster, P. & Röhl, A. Median-joining networks for inferring intraspecific phylogenies. Mol. Biol. Evol. (1999). doi:10.1093/oxfordjournals.molbev.a026036

135. Leigh, J. W. & Bryant, D. POPART: Full-feature software for haplotype network construction. Methods Ecol. Evol. (2015). doi:10.1111/2041-210X.12410

136. Zhang, G. et al. Genetic Associations with Gestational Duration and Spontaneous Preterm Birth. Obstetrical and Gynecological Survey (2018). doi:10.1097/01.ogx.0000530434.15441.45

137. Meunier, J. C. et al. Isolation and structure of the endogenous agonist of opioid receptor-like ORL 1 receptor. Nature (1995). doi:10.1038/377532a0

