## Supplementary Figures 1 and 2 for "Accounting for diverse evolutionary forces reveals the mosaic nature of selection on genomic regions associated with human preterm birth" (from another version of this preprint: version 1, 2019-10-24)

Supplementary Material

Table of Contents:

Supplementary Tables:

1. sPTB lead SNPs and LD SNPs for independent sPTB genomic regions

2. Heatmap z-score results in table format

3. Heatmap p-value results in table format

4. 1KG Individual IDs used for analysis

5. Variant data for analysis

Supplementary Figures:

1. Distribution of matched and unmatched genome wide distributions

2. Sensitivity analysis of diverse evolutionary forces acting on sPTB-associated genomic regions

Supplementary Tables:

**Supplementary Table 1:** All sPTB lead SNPs and LD SNPs tested in pipeline

**Supplementary Table 2:** Heatmap results for zscore in table format

**Supplementary Table 3:** Heatmap results for p-value in table format

**Supplementary Table 4:** 1KG Individual IDs used for analysis

**Supplementary Table 5:** RegulomeDB, GTEx, VEP, and frequency data for significant variants

**Supplementary Figure 1: Distribution of matched and unmatched genome wide distributions for diverse evolutionary measures.**

Each figure below includes sPTB-associated genomic regions (x-axis) that was statistically significant using either a genome wide background (blue dot) or a matched background for the median value of a specific evolutionary measure (red dot) or by both distributions (black dot) (y-axis). Genome wide background distribution (‘Unmatched Bkg, x-axis) is the filled violin plot; all unfilled violin plots are the MAF and LD matched distributions labeled by the lead variant for that sPTB genomic region. Dotted horizontal lines are genome wide unmatched thresholds for one or two-sided p-value of 0.05.

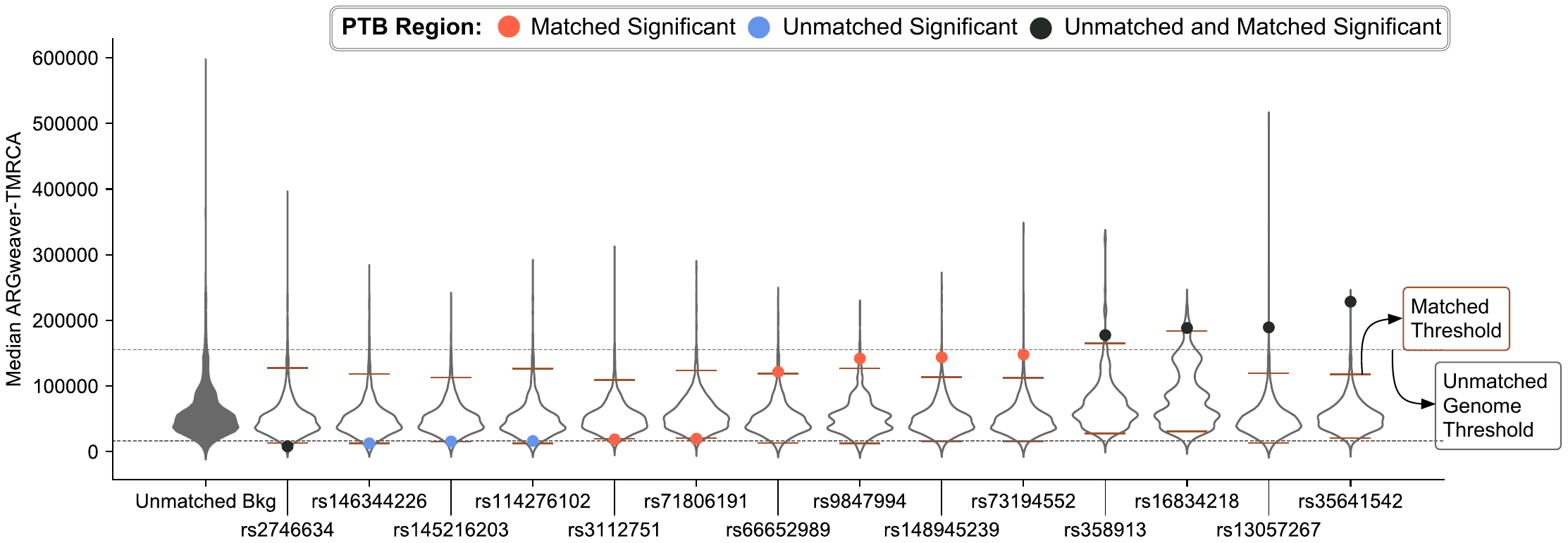

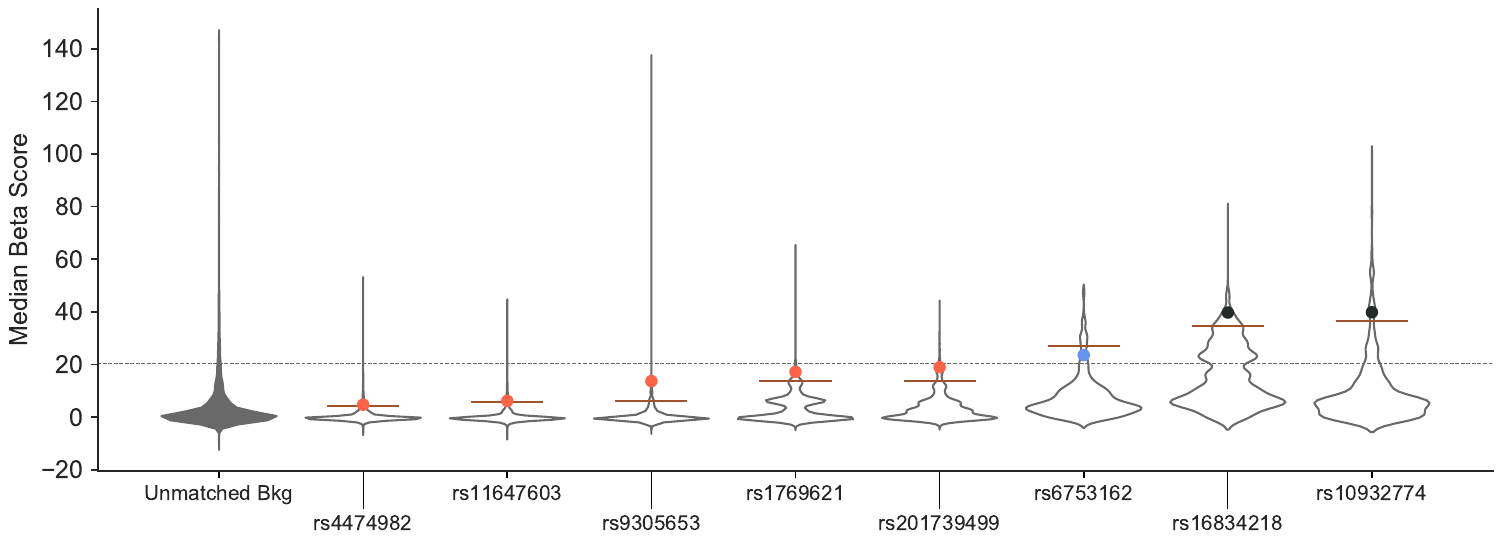

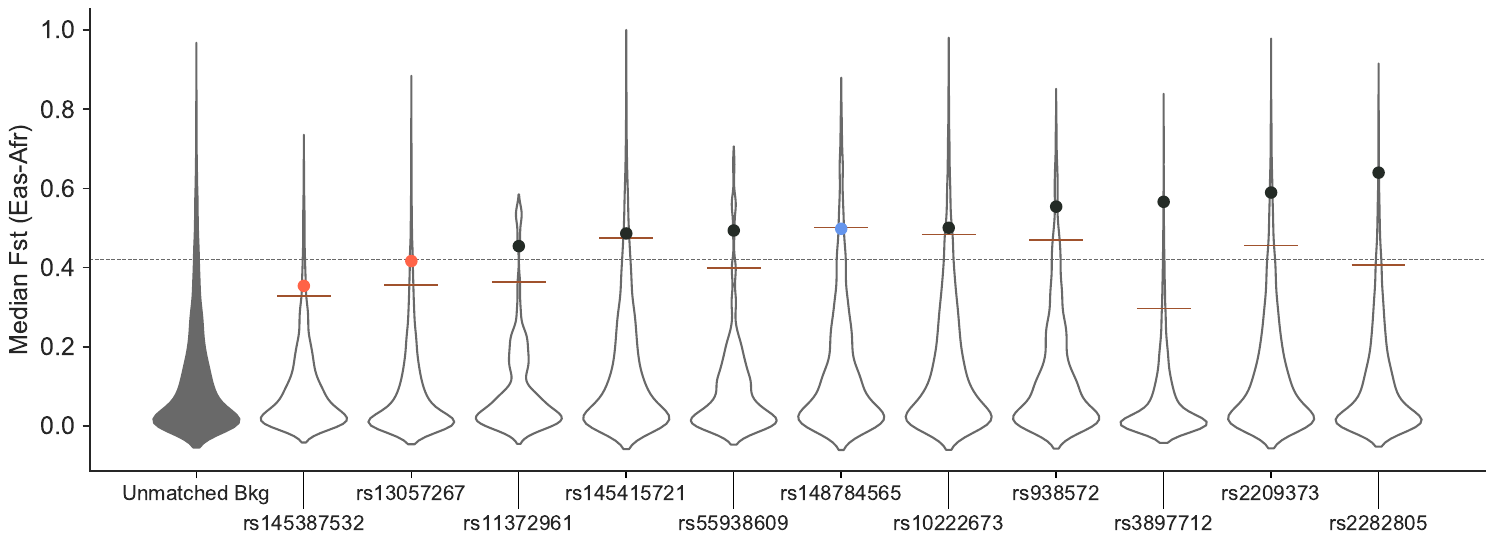

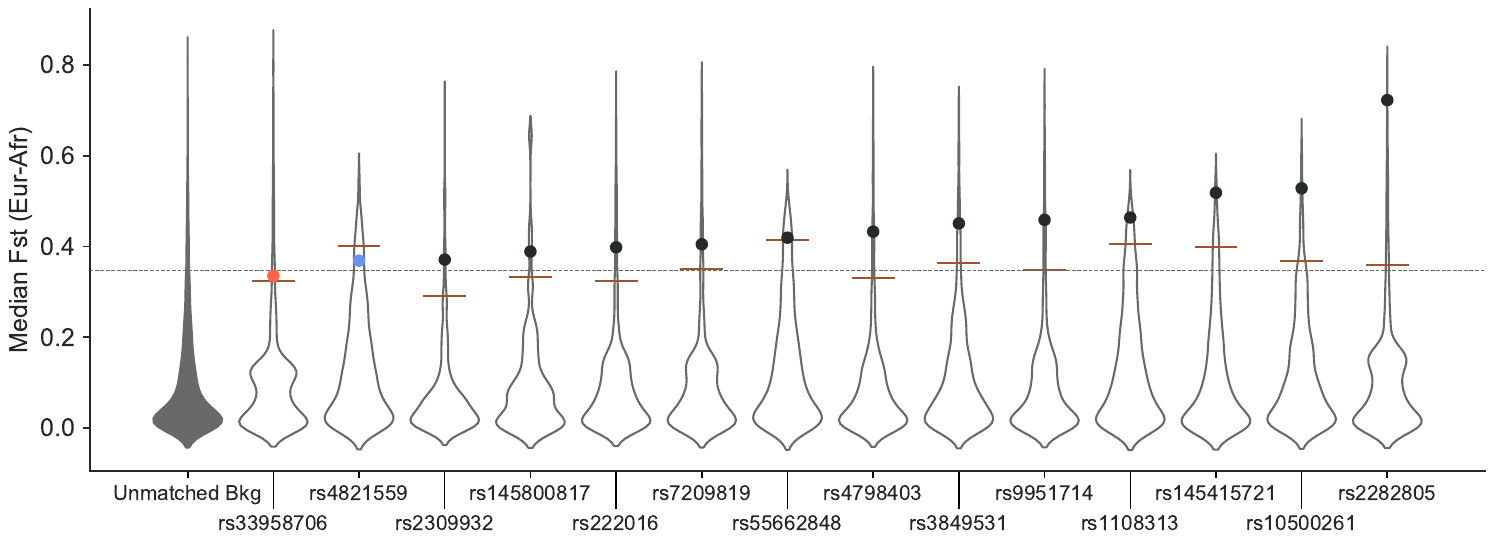

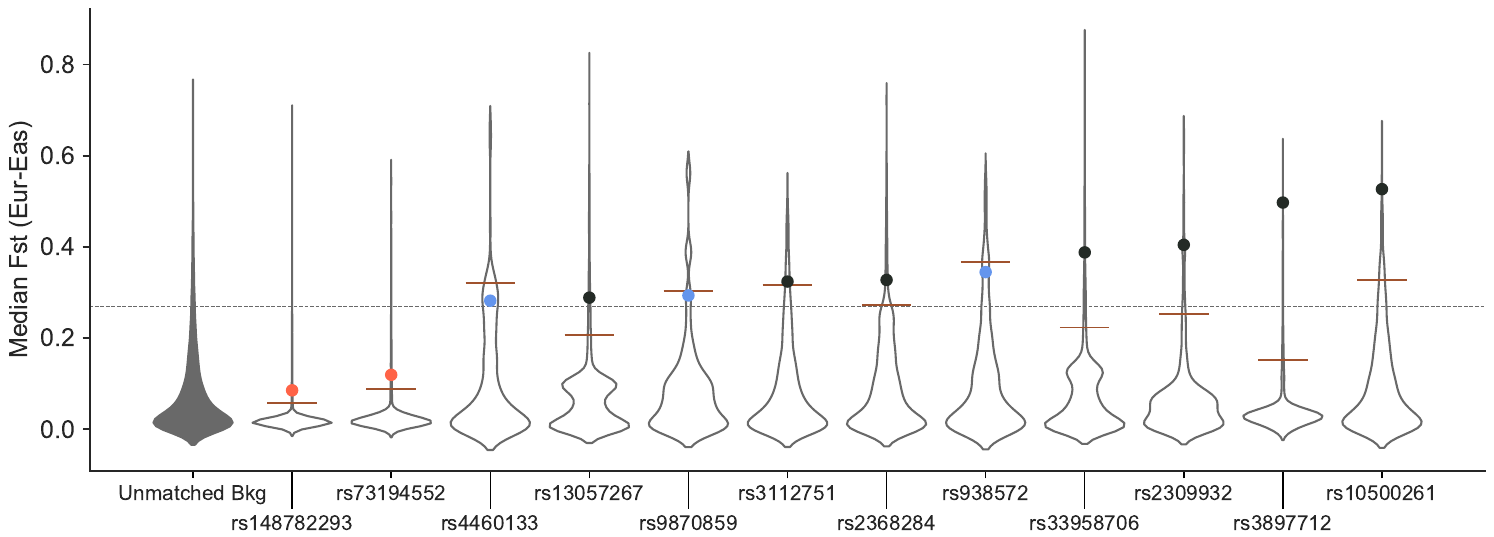

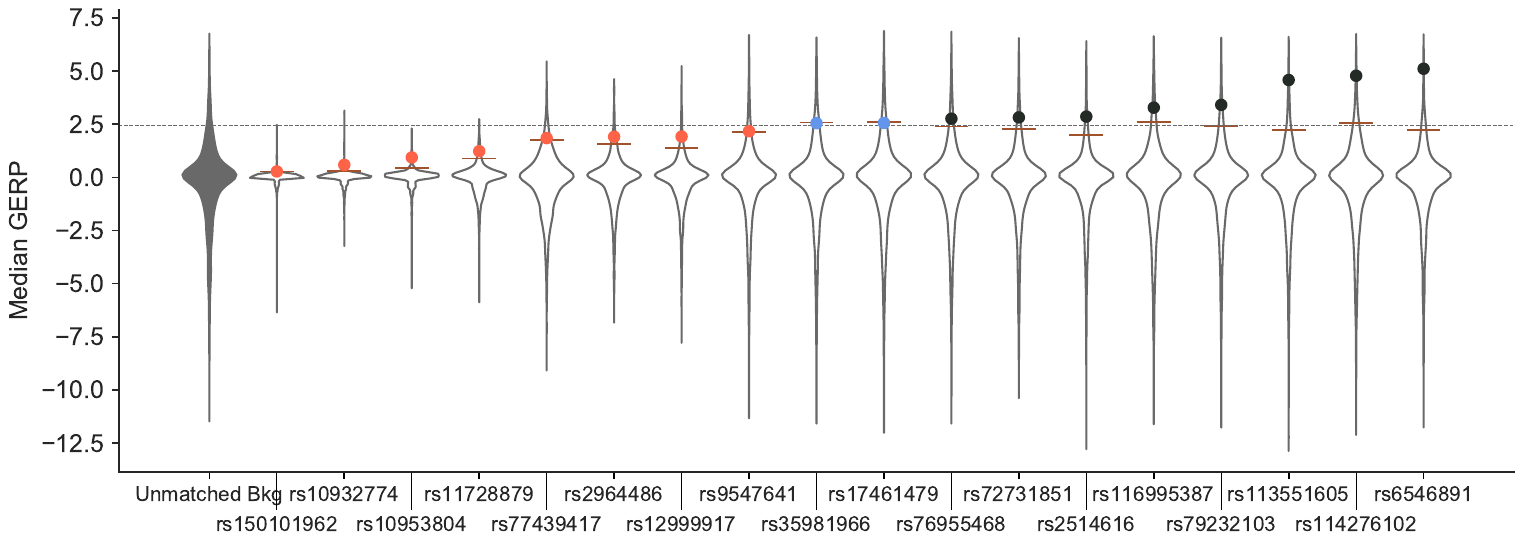

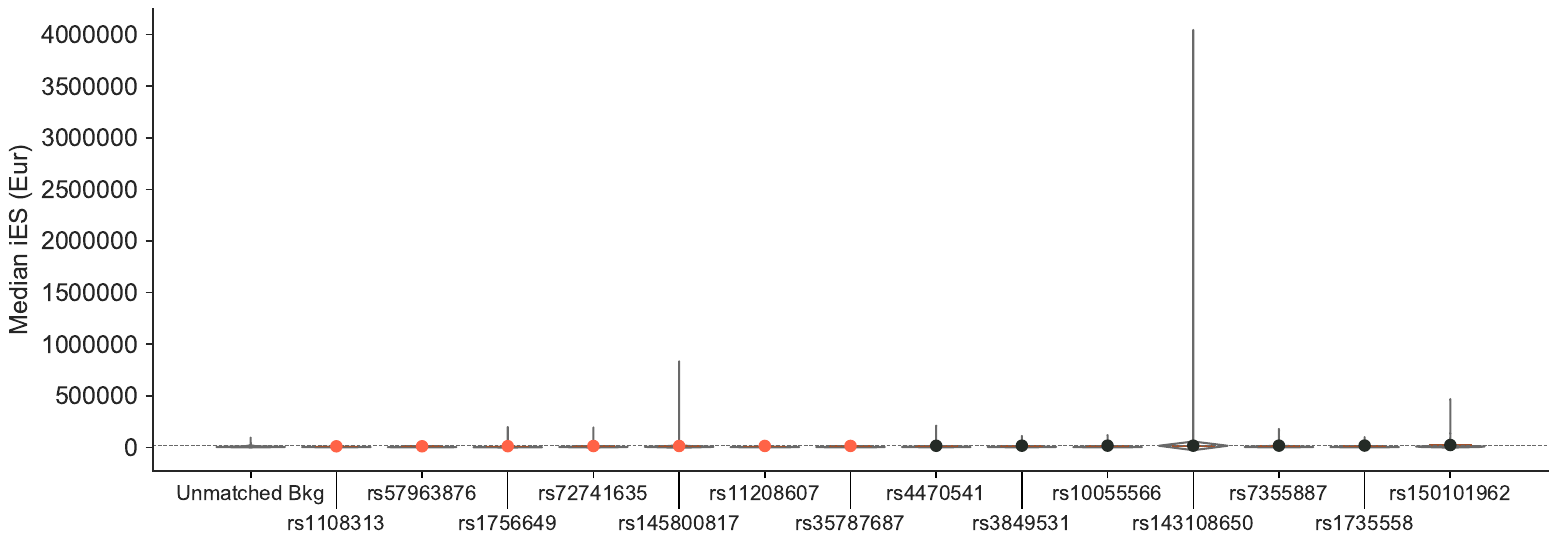

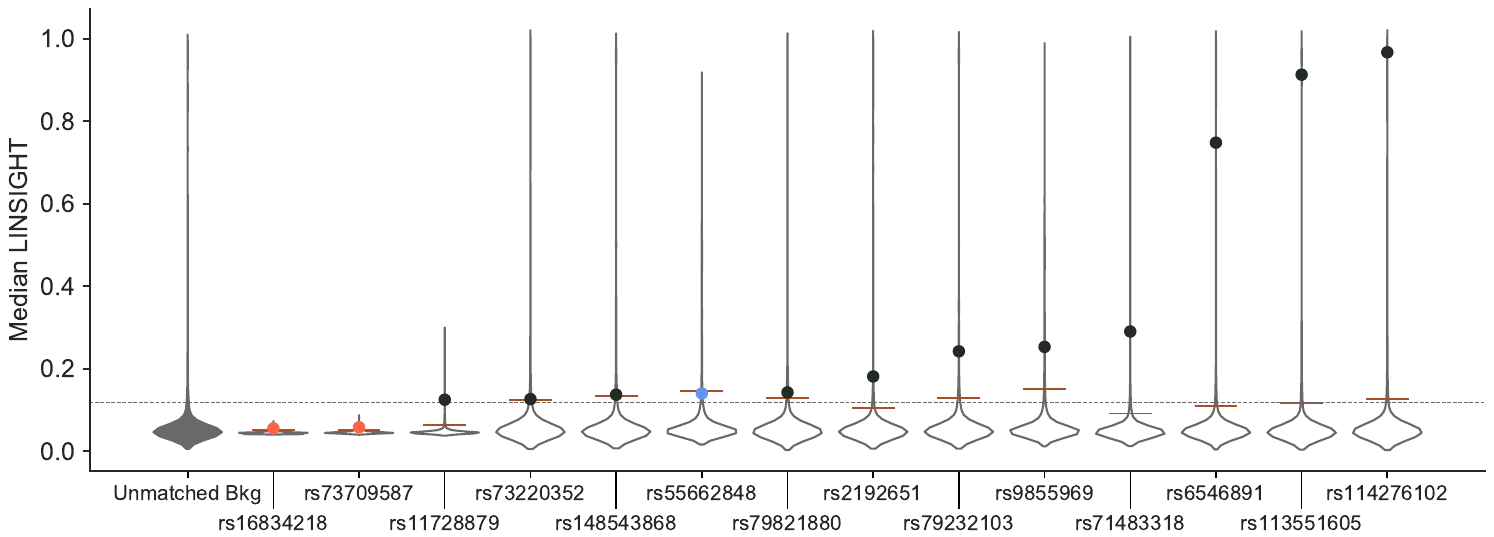

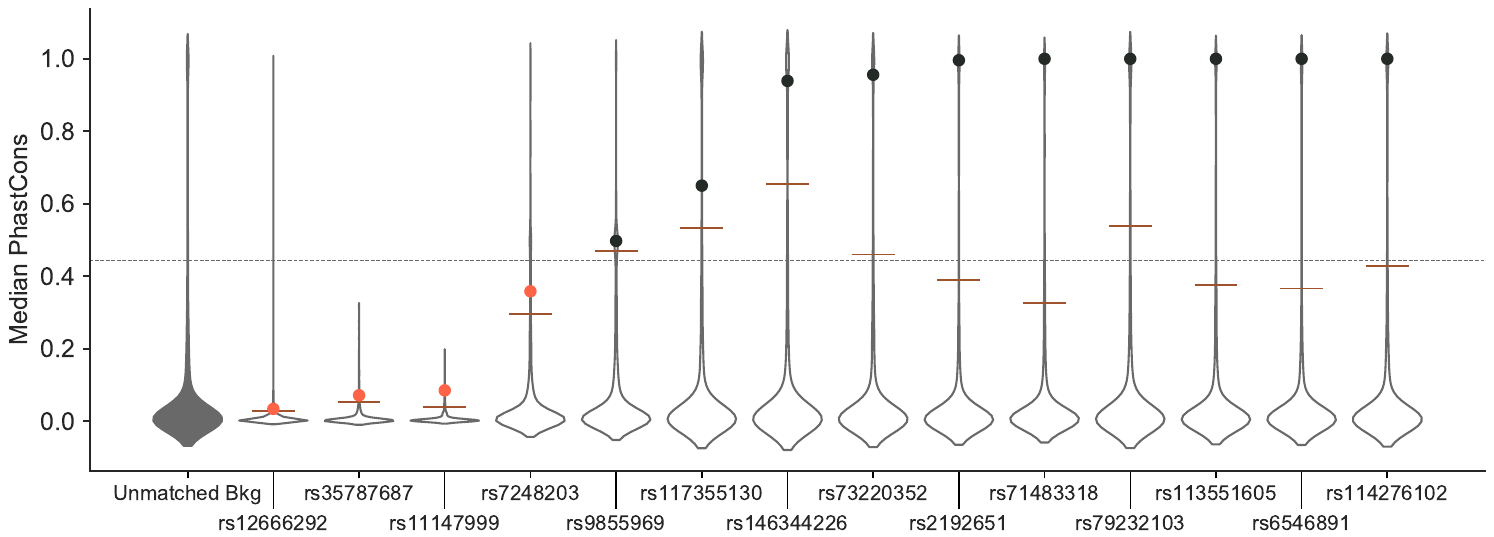

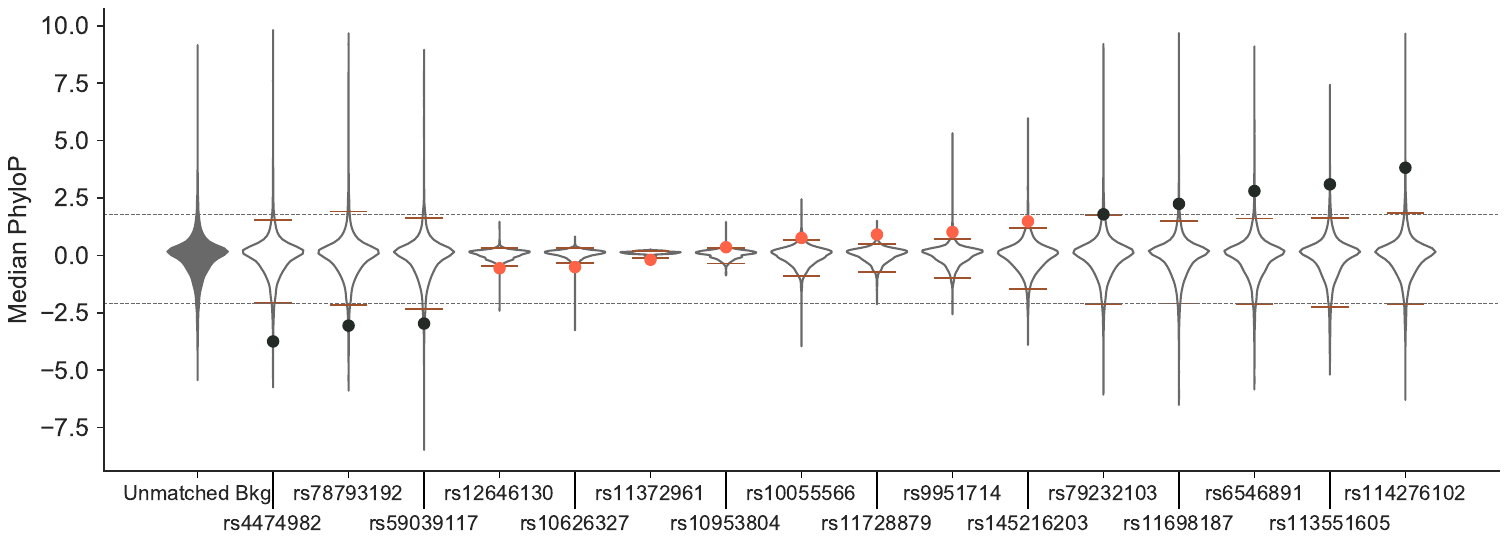

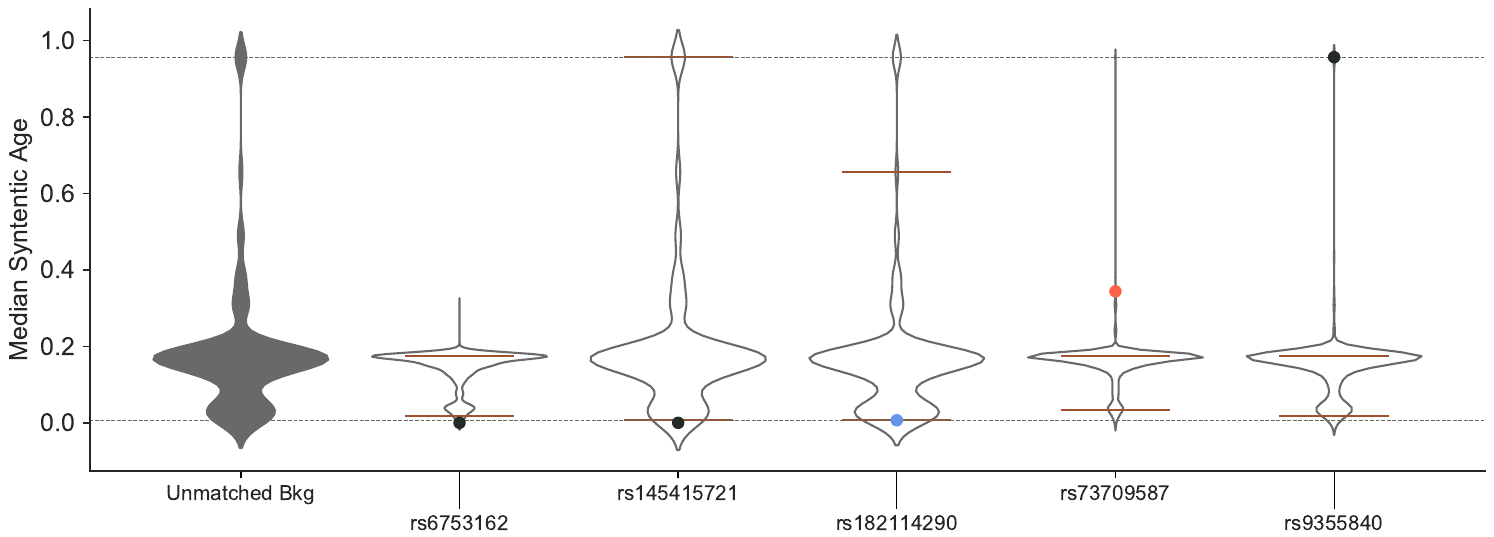

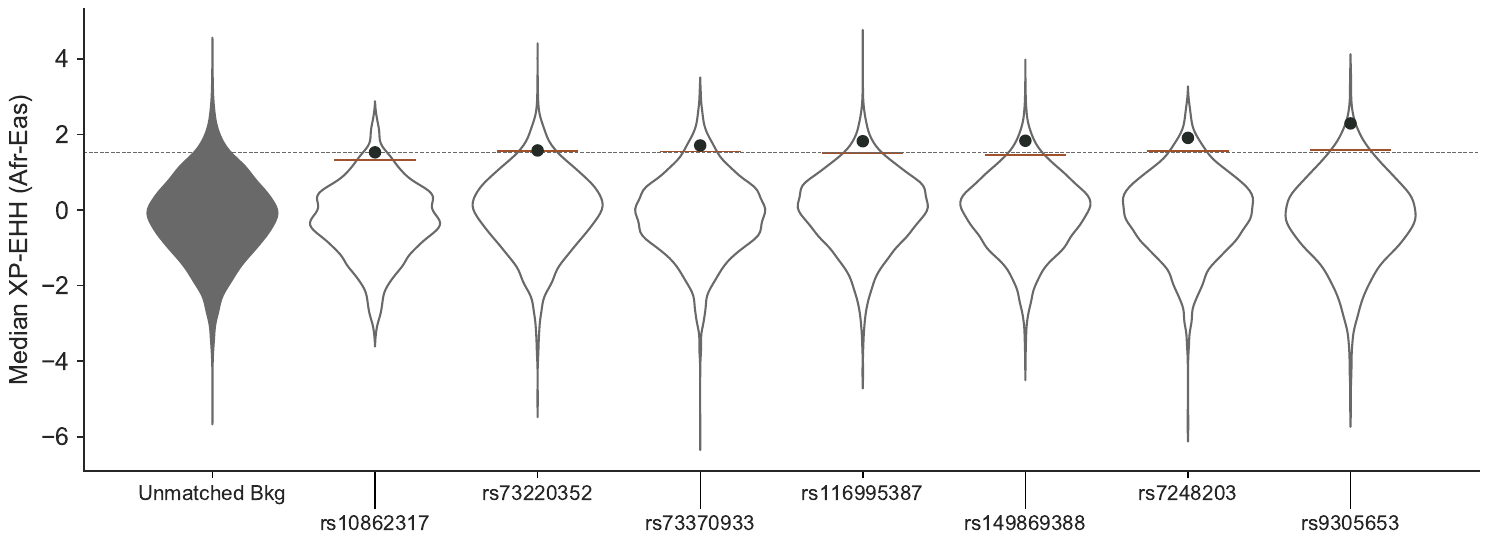

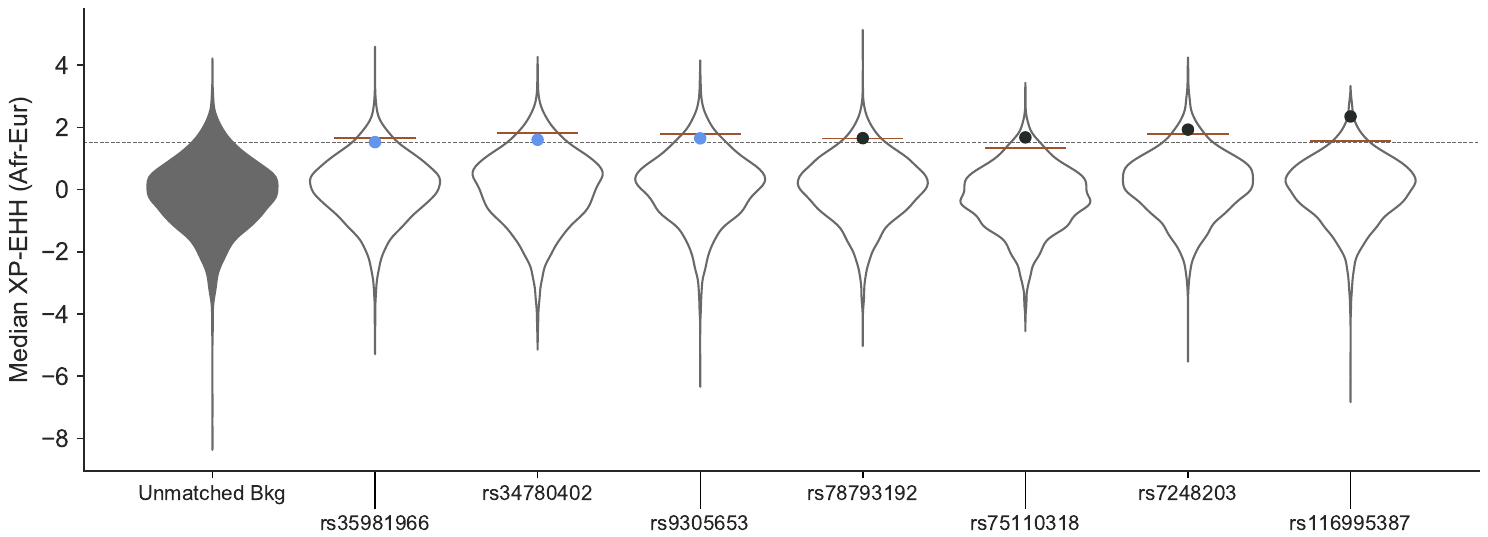

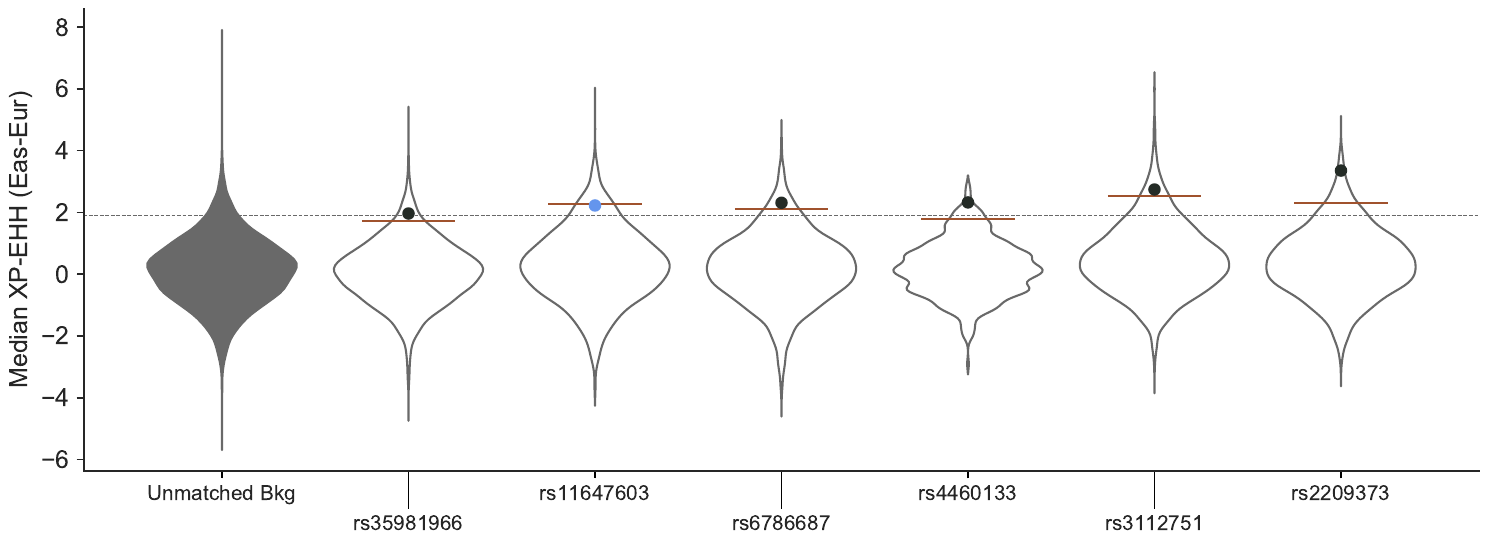

**Supplementary Figure 2: Sensitivity analysis of diverse evolutionary forces acting on sPTB-associated genomic regions**

We tested 215 sPTB-associated genomic regions tested for diverse evolutionary forces (as described in Figure 2 and Methods). Heatmaps show sPTB-associated genomic region (columns) referred to by the lead SNP rsID and evolutionary forces (y-axis). The relative strength (size of square) and direction (color) of each selection measure for each PTB region is presented as a z-score calculated from that region’s background control distribution. Statistical significance was assessed by comparing the median value of the evolutionary measure to the background control distribution to derive an empirical p-value (*p>0.05). We evaluated the pattern of evolutionary forces observed under three different p-value thresholds for the lead variant p-value in each sPTB-associated genomic region: a) p<10E-4, b) 5*10E-5, c) 1*10E-5. Across these thresholds we observed strong signals of diverse evolutionary forces.

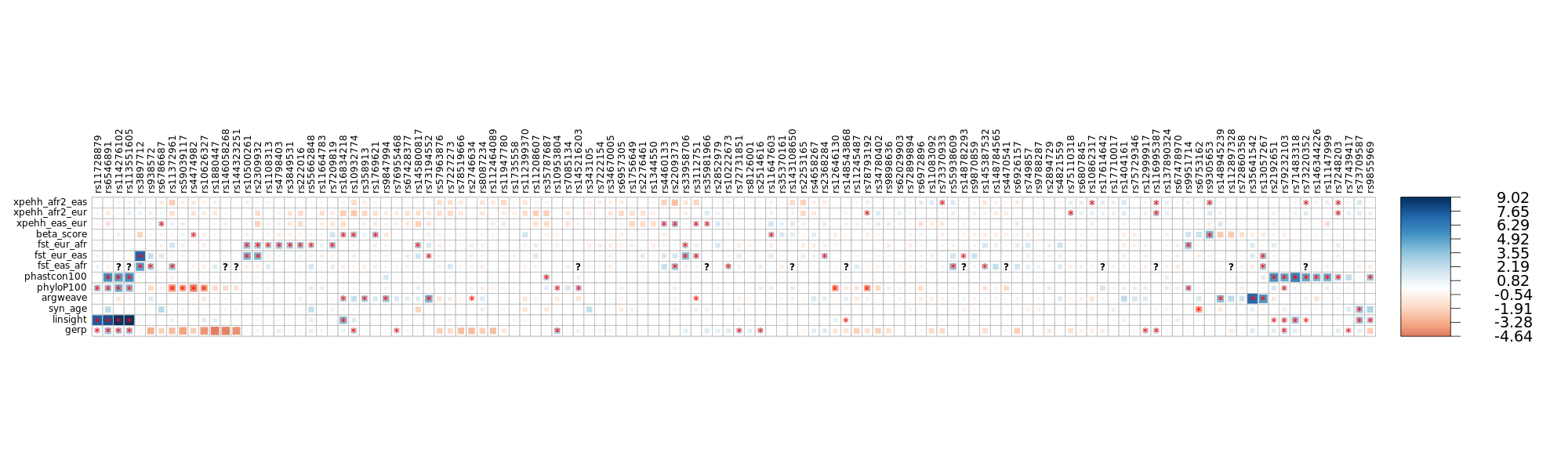
a)

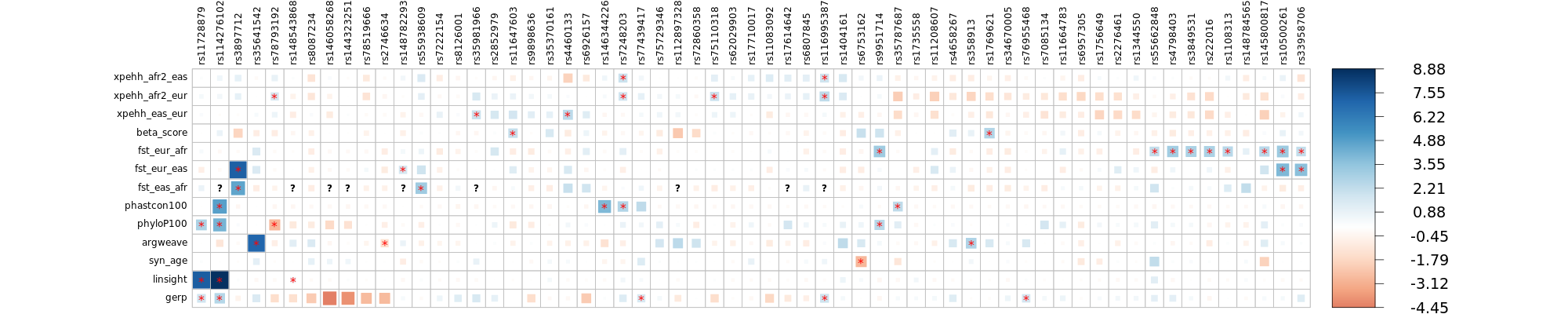
b)

c)

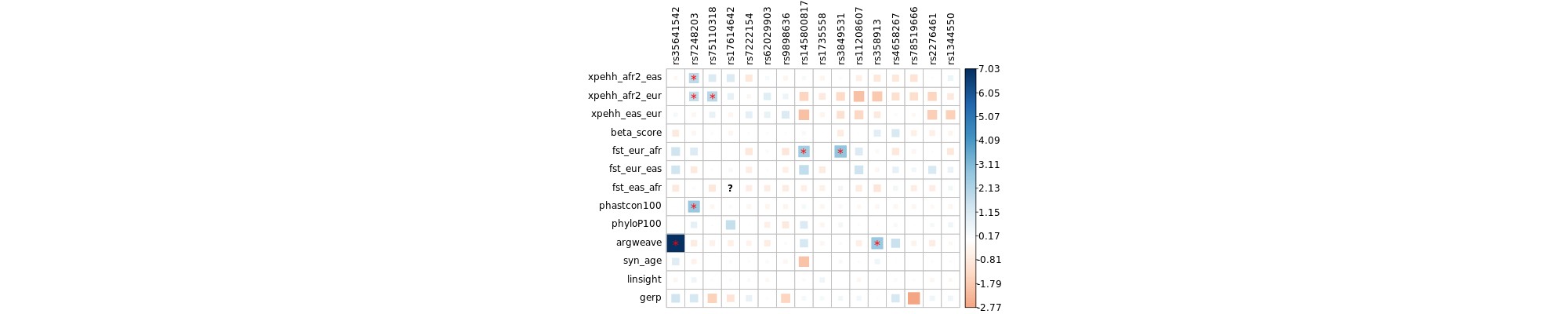
